# Old drugs with new tricks: Efficacy of fluoroquinolones to suppress replication of flaviviruses

**DOI:** 10.1101/2020.03.30.016022

**Authors:** Stacey L. P. Scroggs, Christy C. Andrade, Ramesh Chinnasamy, Sasha R. Azar, Erin E. Schirtzinger, Erin I. Garcia, Jeffrey B. Arterburn, Kathryn A. Hanley, Shannan L. Rossi

## Abstract

Antiviral therapies are urgently needed to treat infections with flaviviruses such as Zika (ZIKV) and dengue (DENV) virus. Repurposing FDA-approved compounds could provide the fastest route to alleviate the burden of flaviviral diseases. In this study, three fluoroquinolones, enoxacin, difloxacin and ciprofloxacin, curtailed replication of flaviviruses ZIKV, DENV, Langat (LGTV) and Modoc (MODV) in HEK-293 cells at low micromolar concentrations. Time-of-addition assays revealed that enoxacin suppressed ZIKV replication when added at 6 hours post-infection, suggesting inhibition of an intermediate step in the virus life cycle, whereas ciprofloxacin and difloxacin had a wider window of efficacy of 2, 6, and 8 hours post-infection for difloxacin and 2 to 8 hours post-infection for ciprofloxacin. The efficacy of enoxacin to suppress ZIKV replication in 5-week-old A129 mice was evaluated in two experiments. First, mice were infected with 1×10^5^ plaque-forming units (pfu) ZIKV FSS13025 (n=20) or PBS (n=11) on day 0 and subsets were treated with enoxacin at 10mg/kg or 15mg/kg or diluent orally twice daily on days 1-5. Treated and control mice did not differ in weight change or virus titer in serum or brain. Mice treated with enoxacin showed a significant, 5-fold decrease in ZIKV titer in testes relative to controls. Second, mice were infected with 1×10^2^ pfu ZIKV (n=13) or PBS (n=13) on day 0 and subsets were treated with 15mg/kg oral enoxacin or diluent twice daily on days 0 (pre-treatment) and 1-5. Mice treated with enoxacin showed a significant, 2.5-fold decrease in ZIKV titer in testes relative to controls, while weight and viral load in the serum, brain, and liver did not differ between treated and control mice. Enoxacin efficacy in cultured murine Sertoli cells was not enhanced compared to efficacy in HEK-293 cells. ZIKV can be sexually transmitted, so reduction of titer in the testes by enoxacin should be further investigated.

**Author Summary:** Flaviviruses such as Zika and dengue virus pose a significant threat to public health worldwide, and there are currently no antiviral therapies to treat any flaviviral infection. Repurposing FDA-approved drugs as anti-flaviviral therapies can accelerate clinical use. We demonstrated that fluoroquinolone antibiotics exhibit anti-flaviviral efficacy, suppressing flavivirus replication in cultured human cells. Additionally, we found that the fluoroquinolone enoxacin suppressed Zika virus replication in mouse testes. While Zika virus is primarily transmitted via mosquitoes, the virus also undergoes sexual transmission. The importance of sexual transmission for the overall epidemiology of the virus remains unclear; nonetheless all routes of potential transmission to pregnant women are of concern as fetal infection *in utero* can have devastating effects. Thus, our data indicate that fluoroquinolones hold promise for treatment of flaviviral infections, particularly infection of the testes by Zika virus, and that this class of drugs warrants further study.

## Introduction

Viruses of the genus flavivirus (family *Flaviviridae*) are major causes of morbidity and mortality worldwide (1–6). Moreover many flaviviruses, such as Zika (ZIKV), dengue (DENV), tick-borne encephalitis (TBEV), West Nile (WNV), and Japanese encephalitis (JEV) virus (7–9) are categorized as emerging pathogens due to rising incidence and expanding geographic range (10,11). Effective antiviral drugs could abate flavivirus transmission and disease burden, but to date no drugs for treatment of flavivirus infections have been brought to market because efforts to develop anti-flaviviral drugs have been unsuccessful (12,13). Most anti-flaviviral drug candidates have stalled at the point of hit-to-lead optimization due to poor drug-like properties (14–16). This history of roadblocks in development of novel drugs suggests that repurposing clinically approved drugs offers the fastest track to clinical treatments of flavivirus infections (17).

The fluoroquinolones are not an immediately obvious choice as anti-flavivirals. The flavivirus genome comprises a single, positive-sense RNA, while fluoroquinolones are primarily known for inhibiting topoisomerases and gyrases in bacterial targets (18,19), neither of which play a role in genome synthesis in positive-sense RNA viruses (20,21). However in the last several years, a multitude of previously unsuspected effects of fluoroquinolones on eukaryotic cell functions have been revealed, including enhancement of RNAi (22–24), inhibition of cellular helicases (25,26), attenuation of cytokines and pro-inflammatory reactive oxygen species (27– 29), and modification of apoptosis (30) and autophagy (31). Furthermore, fluoroquinolones have been shown to suppress hepatitis C virus (HCV, family *Flaviviridae*) replication *in vitro*, possibly by inhibiting the viral helicase (32), but this suppression has not translated into an effective treatment for patients with liver failure due to chronic HCV infection (33). Additionally, fluoroquinolones suppress rhinovirus infection by reducing expression of the viral receptor on cells (34). Recently, Xu and colleagues demonstrated that a high concentration of enoxacin administered to human neuronal progenitor cells (hNPC) and brain organoids prior to and after infection with ZIKV suppressed viral replication and restored normal cellular proliferation, possibly by enhancing RNAi (35).

Here we evaluated the utility of repurposing fluoroquinolones as anti-flavivirals by testing their ability to suppress flavivirus replication in cell culture and a mouse model. This study was initially motivated by our interest in the ability of fluoroquinolones to enhance RNAi, and thus we focused on three fluoroquinolones, enoxacin, ciprofloxacin and difloxacin, that have high, moderate and little impact on RNAi, respectively (23). We found that all three drugs suppressed replication of six flaviviruses in HEK-293 cells at low micromolar concentrations. Enoxacin displayed the lowest EC_50_ values in cell culture and was selected for evaluation in ZIKV-infected A129 mice. Although enoxacin did not mitigate weight loss in ZIKV-infected mice or suppress ZIKV replication in the serum, brain, or liver, the drug did suppress ZIKV replication in the testes.

## Material and Methods

### Viruses

The seven flaviviruses utilized in this study are listed in Table 1. Working stocks of viruses were propagated in Vero cells and viral supernatants were collected either in 1X SPG (2.18 mM sucrose, 38 mM potassium phosphate [monobasic], 72 mM potassium phosphate [dibasic], 60 mM L-glutamic acid) [DENV-1,2 and 4, MODV, LGTV, ZIKV MEX 1-7] for studies in culture or 1X DMEM supplemented with 5% heat inactivated fetal bovine serum (FBS, Atlantica Biologicals, Flowery Branch, GA) and 100μg/mL penicillin/streptomycin (Gibco, Life Technologies, Grand Island, NY) [ZIKV FSS13025] for studies *in vivo*. Supernatants were clarified by centrifugation, aliquoted and stored at −80 °C. Viral titers were determined via serial dilution onto HEK-293 cells followed by immunostaining using methods as previously described (36,37). Briefly, each virus was subjected to serial tenfold dilution and inoculated onto confluent HEK-293 cells in 24-well plates. After two hours of incubation at 37°C with occasional rocking, infected cells were overlaid with 1% methylcellulose in OptiMEM (Gibco, Life Technologies, Grand Island, NY) that had been supplemented with 2% FBS (Gibco, Life Technologies, Grand Island, NY), 2mM L-glutamine (Gibco, Life Technologies, Grand Island, NY), and 0.05 mg/mL gentamycin (Gibco, Life Technologies, Grand Island, NY). Plates were incubated for five days under maintenance conditions, after which cells were fixed with ice cold methanol: acetone (1:1) for 30 minutes. Viral plaques were immunostained using species-specific antibodies and peroxidase-labeled goat anti-mouse secondary antibody (KPL, Gaithersburg, MD) then developed with KPL True Blue Peroxidase Substrate (SeraCare, Milford, MA) and counted to calculate viral titer.

**Table 1.** Passage history for flaviviruses utilized in this study

| **Virus** | **Strain** | **Obtained from** | **Passage history** |
| --- | --- | --- | --- |
| Zika virus (ZIKV) | MEX 1-7 | World Reference Center for Emerging Viruses and Arboviruses (WRCEVA) | C6/36 (x3) |
| Zika virus (ZIKV) | FSS13025 |  | C6/36 (x1), Vero (x1) |
| Dengue virus-1 (DENV-1) | Thailand 160087-1A | Laboratory of Dr. Stephen Whitehead, NIAID, NIH | Vero (x5) |
| Dengue virus-2 (DENV-2) | NGC proto |  | C6/36 (x3), Vero (x2) |
| Dengue virus-4 (rDENV-4) | Dominica p4-3b (36) |  | Vero (x4) |
| Langat virus (LGTV) | E5 (38) | Laboratory of Dr. Alexander Pletnev, NIAID, NIH | Vero (x4) |
| Modoc virus (MODV) | 7/26/61 | WRCEVA | IC suckling mice (x9), Vero (x4) |

### Cells

HEK-293 and murine Sertoli cells were purchased from ATCC (CRL-1573 and CRL-2618, Manassas, VA). Vero cells were obtained from the lab of Stephen Whitehead (NIAID, NIH). HEK-293 cells were maintained at 37°C with 5% CO_2_ in DMEM/F12 medium (Gibco, Life Technologies, Grand Island, NY) supplemented with 10% heat-inactivated FBS (Gibco), 2mM L-glutamine (Gibco), and 0.5% antibiotic-antimycotic (penicillin, streptomycin, and amphotericin B; Gibco). Sertoli cells were maintained at 32°C with 5% CO_2_ in DMEM/F12 (Gibco) supplemented with 10% heat-inactivated FBS, 2 mM L-glutamine, and penicillin/streptomycin (100 units/mL and 100 μg/mL, respectively; Gibco). Vero cells were maintained at 37°C with 5% CO_2_ in DMEM (Gibco) supplemented with 10% heat-inactivated FBS (Gibco). All cell culture efficacy and toxicity experiments were conducted with HEK-293 cells, an interferon competent human cell line that supports flavivirus replication and is often used to evaluate potency and toxicity of potential antivirals (39–45).

### Fluoroquinolone compounds

For each experiment, a fresh working stock of enoxacin (Sigma-Aldrich, E3764, St. Louis, MO), difloxacin (Sigma-Aldrich, D2819, St. Louis, MO), or ciprofloxacin (Corning, 86393-32-0, Manassas, VA) at a concentration of 1.5 mM was sonicated in nanopore water with 3 mM lactic acid (Sigma-Aldrich, L1750, St. Louis, MO) and sterilized *via* passage through a 0.2 μm filter. The compounds were diluted to their final concentrations in cell culture media for assays in cell culture, or nanopore water for *in vivo* treatments.

### Viral replication kinetics in cell culture

To quantify replication kinetics of particular viruses, triplicate 25-cm^2^ flasks of HEK-293 cells were grown to ∼80% confluence, washed with 3mL cell culture media, and infected with a specified virus at a multiplicity of infection (MOI) of 0.05 in 1mL total volume. Cells were incubated at 37°C for 2 hours with occasional rocking. Virus inoculum was then removed and cells washed twice with 3mL of 1x PBS to remove any unadsorbed virus. Six mL of cell culture media was then added to each flask. At time 0, 1mL of cell culture supernatant was removed and SPG was added at a final concentration of 1X. Cell culture supernatants were clarified by centrifugation, aliquoted and stored at −80°C. Samples were collected on days 1 through 8 by removing 1mL of supernatant as described above and 1 mL of cell culture media was added back to the flask. Viral titers were determined in HEK-293 cells as described above. ZIKV was added to this project after the replication kinetics assays were completed, in response to the Public Health Emergency of International Concern declared on February 1^st^, 2016, thus this assay was not conducted with ZIKV.

### Determination of half-maximal effective concentration (EC_50_) against select flaviviruses

To determine the EC_50_ of enoxacin, difloxacin, and ciprofloxacin, monolayers of 80% confluent HEK-293 cells in 24-well plates were infected with either ZIKV, DENV-1, DENV-2, DENV-4, LGTV, or MODV in triplicate at a multiplicity of infection (MOI) of 1. The assay was repeated for all three fluoroquinolones with ZIKV at an MOI of 0.2. The virus was allowed to adsorb for 2 hours at 37 °C after which cells were washed with 1 mL 1x phosphate buffered saline (PBS) to remove unadsorbed virus. Each drug was diluted in a two-fold dilution series in cell culture media, with final concentrations ranging from 150 μM to 4.7 μM, and one mL was added to triplicate treatment wells. Triplicate control wells were treated with cell culture media alone and another set of controls were treated with cell culture media containing 3mM concentration lactic acid, the drug diluent. Infected cells were incubated for five days at normal conditions, after which viral supernatants were collected and viral titers were determined as described above.

As enoxacin was found to suppress ZIKV in the mouse testes, enoxacin potency was evaluated in one testicular cell line (murine Sertoli cells) and compared to the potency in HEK-293 cells. The EC_50_ methods described above were repeated for ZIKV MEX 1-7 in Sertoli cells and HEK-293 cells, both incubated at 32 °C to control for potential differences in enoxacin activity at the lower temperature required for Sertoli cell viability. For both cell types, two MOIs were tested, 0.1 and 1.0, and virus was collected at two time points, 2 days post infection (p.i.) and 5 days p.i. Viral titers were determined in HEK-293 cells as described above.

### Determination of half-maximal cytotoxic concentration (CC_50_) of fluoroquinolones

To determine the toxicity of enoxacin, difloxacin, and ciprofloxacin, HEK-293 cells were grown in 96-well plates until confluent at which time the media was removed. Each filter-sterilized fluoroquinolone was diluted two-fold, starting at 500 μM, and added to wells in triplicate at a total volume of 100 μL. Control wells were treated with 100 μL of cell culture media containing 3 mM lactic acid. Plates were incubated at normal conditions for five days, after which the media was removed and 110 μL of 10% resazurin dye (Millipore Sigma, St. Louis, MO) diluted in cell culture media was added to each well. After two hours incubation, absorbance was measured on a plate reader at 600 nm and normalized to the mean absorbance of the control wells.

### Time-of-addition assays

Time-of-addition assays were conducted to gain insight into the potential mechanism of action of each drug against ZIKV (46–50). All assays were conducted in triplicate; MOI and drug concentration were varied in order to enhance statistical power to discern time-specific effects. First, the impact of enoxacin, ciprofloxacin and difloxacin were tested at 24.4 μM, 116.1 μM, and 35.9 μM, respectively, against ZIKV at an MOI of 0.2. These drug concentrations represent the EC_50_ values determined in HEK-293 cells infected with ZIKV at an MOI of 0.2. Next, the time-of-addition assays were conducted using 18.1 μM enoxacin, 56.8 μM ciprofloxacin and 25.4 μM difloxacin against ZIKV at an MOI of 1.0. These drug concentrations represent the EC_50_ values determined in HEK-293 cells with ZIKV at an MOI of 1.0. Finally, a third assay was conducted using 25.0 μM ciprofloxacin and 50.0 μM difloxacin against ZIKV at an MOI of 1.0. These concentrations were chosen to ameliorate suppression of ZIKV by ciprofloxacin and increase suppression by difloxacin.

Eight timepoints were evaluated during all time-of-addition assays: two hours prior to infection, at the time of infection (drug mixed with ZIKV), 2, 4, 6, 8, 12, and 18 hours p.i. These time points capture flavivirus binding and entry (−2 and 0 hrs p.i.), translation (2 and 4 hrs p.i.), genome replication (6 and 8 hrs p.i.), and virion assembly and budding (12 and 18 hrs p.i.) (51). At time zero, monolayers of 80% confluent HEK-293 cells in 24-well plates were infected with ZIKV MEX 1-7 at MOI 0.2 or 1. After two hours of incubation, the virus was removed from all wells, the cells were washed with 1 mL of 1x PBS, and 1 mL of media per well was replaced. At each time point, media was removed from designated triplicate wells, cells were washed with 1x PBS, and 1 mL of drug at the specified concentration was added. Dilution of fluoroquinolones to final concentration occurred at the time of treatment. For the wells treated at −2 hrs p.i., the drug was removed and replaced with virus at the time of infection (time 0) then, after 2hrs incubation, the virus was removed, the wells were washed with 1xPBS, and media was added to the wells. For the wells treated at infection (0 hr p.i.), the media was removed and replaced with ZIKV diluted in the drug at the time of infection. After 2 hrs incubation the virus and drug were both removed, the wells were washed, and media was added. For the post-infection time points, the drug was added to the wells at the specified time points and remained in the wells until 24 hrs p.i. It should be noted that the half-life of enoxacin is 1.75 hrs of ciprofloxacin and difloxacin is 3 hours (52–54). Control wells infected with ZIKV were washed two hours p.i. and treated with 1 mL of media per well. At 24 hours p.i. all the viral supernatants were collected, clarified, and stored as described above.

### Determination of *in vivo* efficacy of enoxacin

The impact of enoxacin, the fluoroquinolone with the lowest EC_50_, on ZIKV infection of A129 mice was tested. Mice were infected at 5 weeks of age because our previous work showed that in this age group ZIKV infection caused sustained weight loss that did not require euthanasia until 8 days p.i., ensuring that the majority of mice would survive a 5-day trial (55). Mice were housed in sterile caging in colonies at the University of Texas Medical Branch, an AALAS-accredited facility, and research was conducted in accordance with UTMB policy under IACUC Protocol #1708051.

The efficacy of enoxacin to suppress ZIKV replication *in vivo* was tested in two separate experiments (Fig. 1). An *a priori* power analysis was used to determine the minimum number of mice required to achieve 80% power to detect a difference of 0.3 log (i.e. 50%) decrease in viral replication of the serum of 5-week-old-A129 mice. In experiment 1, we tested two concentrations of enoxacin at 10 mg/kg and at 15 mg/kg in A129 mice infected with 1×10^5^ pfu ZIKV, the viral dose used to in our previous work to characterize ZIKV infection in 5-week-old A129 mice (55). The two concentrations, 10 mg/kg and 15 mg/kg, were selected because in an average sized mouse (20 g) these doses correspond to peak serum concentrations (6.2 μM and 9.4 μM) that are comparable to the peak serum concentrations achieved in humans receiving a typical clinical dose, wherein 200 mg and 400 mg oral dosages result in peak serum concentrations of 5.0 μM and 11.2 μM, respectively (56,57). In experiment 2, we tested the impacts of a lower dose of virus (1×10^2^ pfu) and a pre-infection treatment of enoxacin on ZIKV infection in mice. The experiment was limited to a single concentration of enoxacin, 15 mg/kg, in order to utilize the minimum number of mice.

**Fig 1.**
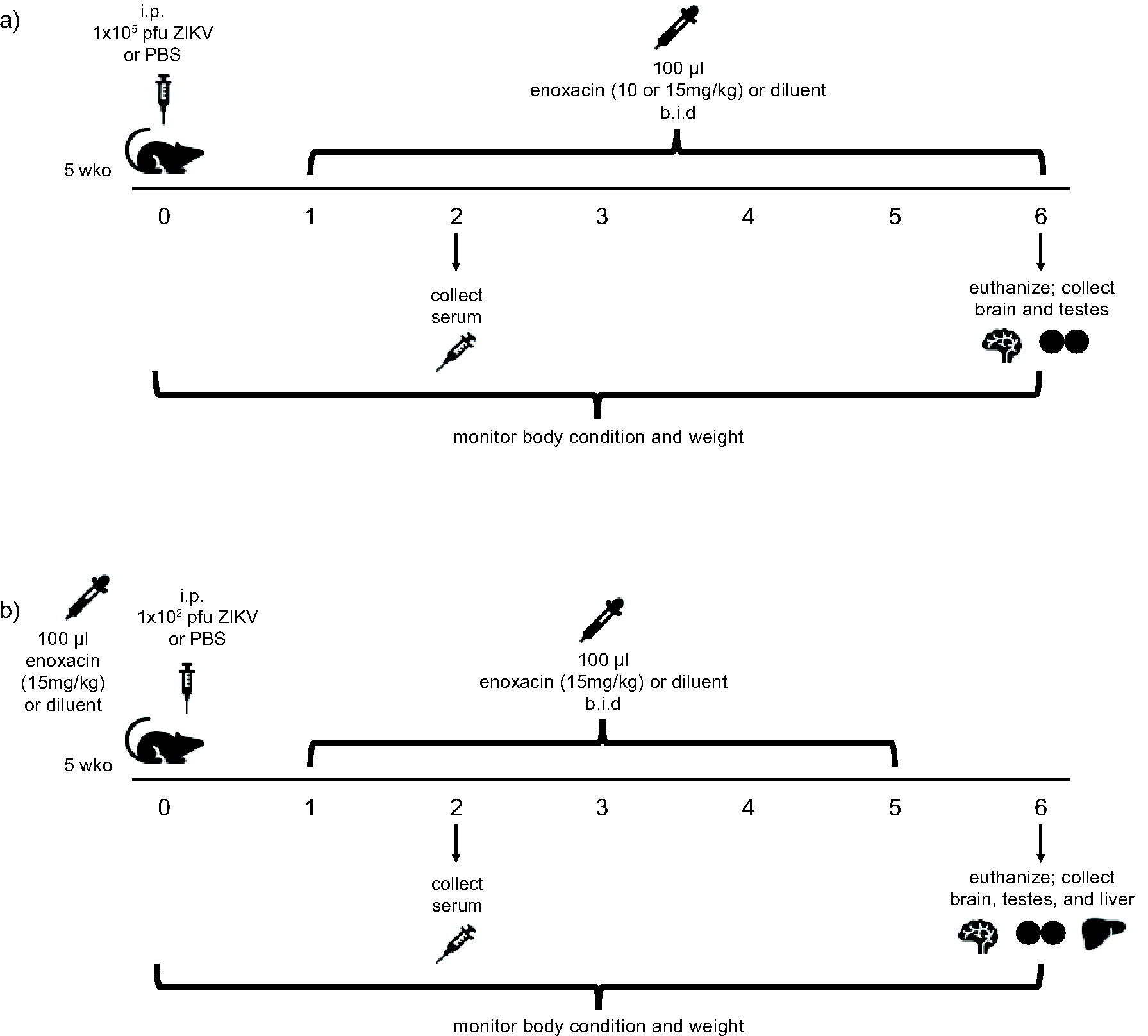
Experimental design. a) In experiment 1, 5-week-old A129 mice were injected with ZIKV (1×10^5^ pfu) or PBS then treated orally with enoxacin (10 or 15 mg/kg) or drug diluent twice daily on days 1-6. b) In experiment 2, 5-week-old A129 mice were pre-treated with enoxacin (15 mg/kg) or diluent 8 hours before injection with ZIKV (1×10^2^ pfu) or PBS and then were treated orally with enoxacin (15 mg/kg) or diluent twice daily on days 1-5 p.i.

#### Experiment 1 (Fig. 1a)

Mice were intradermally injected on day 0 with 1×10^5^ pfu ZIKV FSS13025 diluted in 1x PBS (n=19) or with 1x PBS as a control (n=11) in a total volume of 100 μL and subsets of infected and uninfected mice were treated with oral enoxacin or drug diluent (3 mM lactic acid) (Table 6) twice daily on days 1-6 p.i. Weight and body condition were recorded twice daily. Two days p.i., 70 μL of blood was collected from the retro-orbital sinus, clarified by centrifugation (5 minutes at 3380 x g), and serum was stored at −80 °C. Six days p.i., mice were euthanized and brain and testes were collected. Each tissue, along with a sterile steel ball, were placed into a 2 mL Eppendorf tube containing 500 μL DMEM supplemented with 2% FBS and penicillin/streptomycin and homogenized in a Qiagen TissueLyser II shaking at 26 pulses/second for 5 minutes. Homogenates were clarified by centrifugation at 3380 x g for 5 minutes and stored at −80 °C. Viral titers from serum and tissues were determined in Vero cells in 12 well plates essentially as described above (55).

#### Experiment 2 (Fig. 1b)

Mice received a pre-treatment of 15 mg/kg enoxacin (n= 14) or drug diluent (n = 13) and were intradermally injected with 1×10^2^ pfu ZIKV FSS13025 or 1x PBS 8 hours later as specified in Table 2. Subsets of infected and uninfected mice were treated with oral enoxacin or lactic acid diluent twice daily on days 1-5. Weight and body condition were recorded daily. Serum, brain and liver were collected, and viral titers determined as described above.

**Table 2.** Number of ZIKV-infected and control mice treated with enoxacin or drug diluent

|  | Experiment 1 |  | Experiment 2 |  |
| --- | --- | --- | --- | --- |
| | $1 \times 10^5$ pfu ZIKV | PBS Control | $1 \times 10^2$ pfu ZIKV | PBS Control |
| Drug diluent | 7 (4 female, 3 male) | NA | 7 (3 female, 4 male) | 6 (3 female, 3 male) |
| Enoxacin (10 mg/kg) | 7 (3 female, 4 male) | 6 (2 female, 4 male) | NA | NA |
| Enoxacin (15 mg/kg) | 6 (0 female, 6 male) | 5 (3 female, 2 male) | 6 (3 female, 3 male) | 7 (5 female, 2 male) |

### Statistical analysis

EC_50_ values were calculated using nonlinear regression of inhibition dose response for log drug concentration and viral titer and CC_50_ values were calculated using nonlinear regression of inhibition dose response for log drug concentration and cell viability in GraphPad Prism (version 5 for Mac OS X, GraphPad Software, La Jolla California USA). The selectivity index for each fluoroquinolone and virus combination was calculated by dividing the CC_50_ by the EC_50_ values. Mean viral titers at each time point from the time-of-addition assays were tested for normality using the Shapiro-Wilk test then analyzed using ANOVAs. If the overall ANOVA was significant, pairwise t-tests with a Bonferroni correction were used to detect pairwise differences. Viral titers were first log-transformed then mean viral titers from mice were assessed for normality using the Shapiro-Wilk test and compared using ANOVAs or t-tests as appropriate, and differences in ZIKV replication in Sertoli cells and HEK-293 cells at 32 °C were evaluated using general linear models in R (58).

## Results

### Flavivirus replication curves in cultured human cells

Replication curves for DENV-1, DENV-2, DENV-4, LGTV, and MODV in HEK-293 cells are shown in Fig. S1. Following infection at MOI 0.05, DENV-1 and DENV-4 titer rose steadily through day 8 p.i., the last day of sampling, while DENV-2 peaked on day 7, LGTV peaked on day 3 and MODV plateaued around 4 days p.i. On day 5 p.i. the viral titer for all 5 viruses ranged from 5.3 to 6.7 log10 PFU/mL. In light of the variation in replication dynamics among the different viruses, subsequent evaluations of drug potency were conducted at a higher MOI (MOI:1) on day 5 post-infection to capture the plateau of viral titer.

### Fluoroquinolones suppress flavivirus replication in cultured human cells

As shown in Table 3 and Fig. S2, the EC_50_ values of enoxacin, ciprofloxacin and difloxacin against DENV, ZIKV, LGTV and MODV in HEK-293 cells at 37 ° C were all in the range of 4.3-56.8 µM, except for the EC_50_ of difloxacin for MODV, for which the EC_50_ value was greater than the limit of detection in our assay (Table 3). CC_50_ values for enoxacin, ciprofloxacin, and difloxacin were all substantially higher than the EC_50_ values (Table 3). In general, difloxacin yielded the highest selectivity indices across viruses tested.

**Table 3.** Potency and toxicity of enoxacin, difloxacin, and ciprofloxacin against flaviviruses in HEK-293 cells

| Drug | $\text{CC}_{50} \mu\text{M}$<br>(95% CI) | Virus | $\text{EC}_{50} \mu\text{M}$<br>(95% CI) | Selectivity<br>Index<sup>a</sup> |
| --- | --- | --- | --- | --- |
| Enoxacin | 537.8 (430.1-700.0) | ZIKV (MOI: 0.2) | 24.4 (17.3-34.1) | 22.0 |
|  |  | ZIKV (MOI: 1.0) | 18.1 (14.6-22.4) | 29.7 |
|  |  | DENV-1 | 6.6 (6.0-7.3) | 81.5 |
|  |  | DENV-2 | 4.7 (3.5-6.2) | 114.4 |
|  |  | DENV-4 | 7.6 (7.1-8.2) | 70.8 |
|  |  | LGTV | 4.3 (2.8-6.5) | 125.1 |
|  |  | MODV | 14.6 (7.4-29.0) | 36.8 |
| Difloxacin | 1479.0 (1087.0-2272.0) | ZIKV (MOI: 0.2) | 35.9 (19.0-67.5) | 41.2 |
|  |  | ZIKV (MOI: 1.0) | 25.4 (20.8-30.9) | 58.2 |
|  |  | DENV-1 | 10.9 (9.2-12.9) | 135.7 |
|  |  | DENV-2 | 5.7 (4.8-6.9) | 259.5 |
|  |  | DENV-4 | 10.1 (9.1-11.3) | 146.4 |
|  |  | LGTV | 8.2 (6.3-10.6) | 180.4 |
|  |  | MODV | >150 | n.d. |
| Ciprofloxacin | 759.6 (649.3-912.9) | ZIKV (MOI: 0.2) | 116.1(68.9-179.0) | 6.5 |
|  |  | ZIKV (MOI: 1.0) | 56.8 (39.6-81.5) | 13.4 |
|  |  | DENV-1 | 27.8 (22.1-34.9) | 27.3 |
|  |  | DENV-2 | 8.0 (5.0-12.9) | 95.0 |
|  |  | DENV-4 | 19.6 (16.5-23.2) | 38.8 |
|  |  | LGTV | 7.4 (3.9-14.0) | 102.6 |
|  |  | MODV | 11.2 (3.8-32.6) | 67.8 |

a: Selectivity Index: $\text{CC}_{50}$ divided by $\text{EC}_{50}$
n.d.: not determined

### Fluoroquinolone suppression of different life cycle stages of ZIKV

Three sets of time-of-addition assays were used to discern the viral life stage(s) inhibited by each of the three fluoroquinolones. First, monolayers of HEK-293 cells were infected at MOI 0.2 and treated with each of the three drugs at the EC_50_ value determined using MOI: 0.2. Next monolayers of HEK-293 cells were infected at MOI 1.0 and treated with each of the three drugs at the EC_50_ value determined using MOI: 1.0. In this second set of assays, ciprofloxacin suppressed virus replication below the level of detection and difloxacin had little effect (Fig. S3), so these assays were run again with ciprofloxacin at half its EC_50_ and difloxacin at twice its EC_50_.

While significance varied somewhat among the assays, the overall patterns were consistent. As seen in Fig. 2a and 2b, enoxacin suppressed virus replication most strongly when added at 2, 4 or 6 hours p.i. Virus replication was suppressed by about 25% by 18.1 μM enoxacin and 50% by 24.4 μM enoxacin. At both concentrations, differences between viral titers by time of treatment were significant. Pairwise t-tests indicated that enoxacin treatment at 2, 4, and 6 hours p.i. significantly reduced ZIKV titer compared to the media control (pairwise t-test P <0.05; full statistics in Table S1).

**Fig 2.**
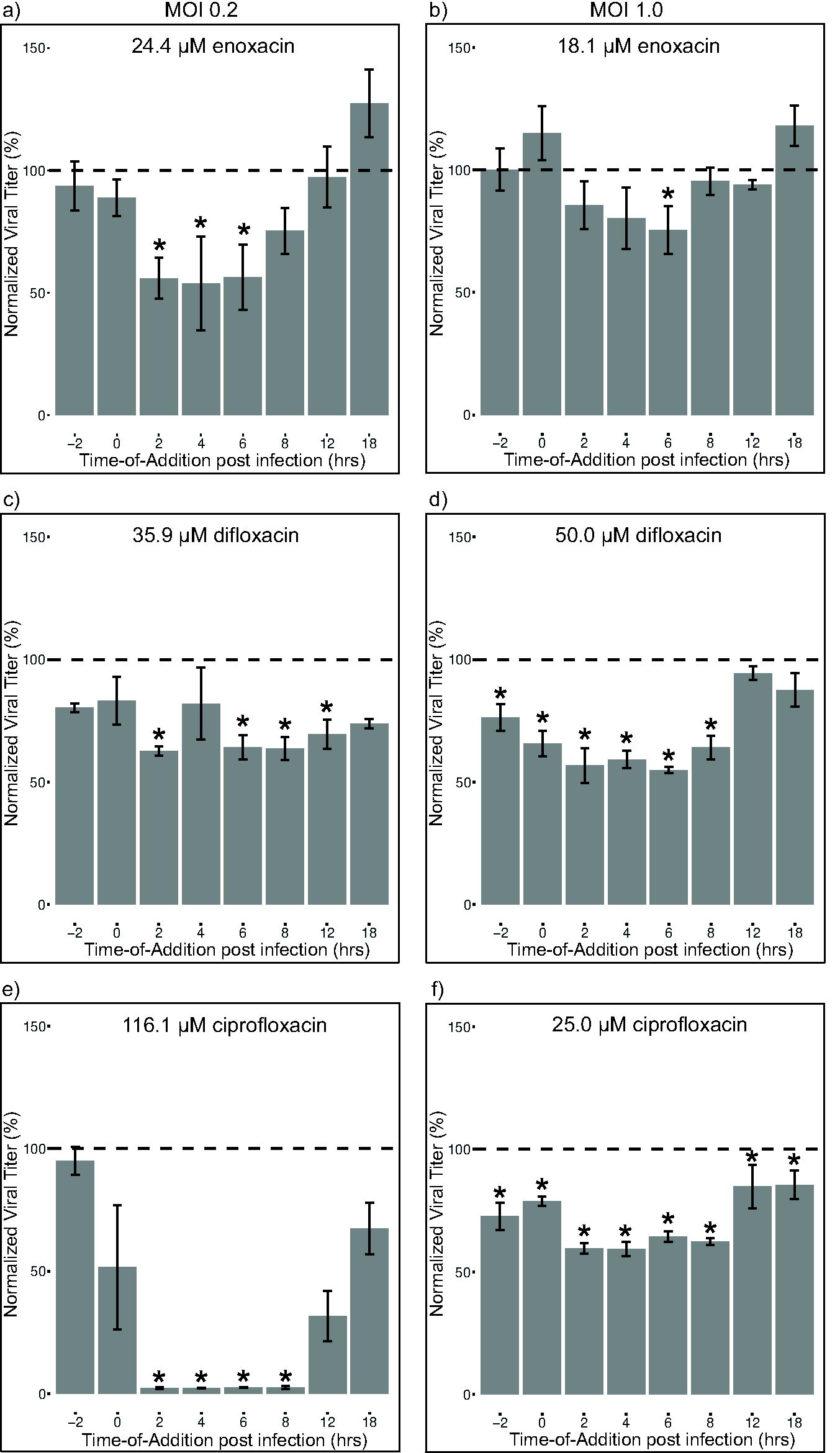
Enoxacin suppresses intermediate life cycle stages of ZIKV while difloxacin and ciprofloxacin suppress early and intermediate life cycle stages of ZIKV. Results of time-of-addition assays of each of three fluoroquinolones against ZIKV at designated drug concentrations and virus MOIs (see text for justification of drug concentration and MOI pairings): for enoxacin (a,b), difloxacin (c,d), and ciprofloxacin (e,f). Viral titers (n = 3 replicates per drug per time point) for each time point were normalized to the average viral titer with media treatment and reported as average percent (titer at time point/average media titer*100). Differences in mean viral titers (log_10_ pfu/mL) were detected with ANOVA and pairwise t-tests; full pairwise statistics in Table S1. * P < 0.05 compared to media control.

Difloxacin at 35.9 μM suppressed virus replication when added 2, 6, 8, 12, and 18 hrs p.i. (Fig. 2c) while 50.0μM difloxacin suppressed virus replication when added 2 hours before infection, at the time of infection, 2, 4, 6, or 8 hours p.i. (Fig. 2d), as detected by pairwise t-tests (full statistics in Table S1). At most, ZIKV replication was suppressed 37% by 35.9 μM difloxacin and 50% by 50.0 μM difloxacin. At 25.4 μM difloxacin, the EC_50_ of this drug against ZIKV at MOI 0.2 (Table 3) no difference in viral titer was detected by time (Fig. S3a, Table S1).

As seen in Fig. 2e and 2f and Fig. S3b, ciprofloxacin most strongly and consistently suppressed virus replication when added 2, 4, 6, or 8 hours p.i. At these time points, virus was suppressed an average of 40% by 25.0 μM ciprofloxacin, 71% by 56.8 μM ciprofloxacin, and below the level of detection by 116.1 μM ciprofloxacin. The differences in viral titers by time of treatment were significant for all three concentrations of ciprofloxacin tested (Table S1).

Pairwise comparisons revealed that 25.0 μM ciprofloxacin added 2 hours before infection, at the time of infection, and up until 18 hours p.i. significantly reduced ZIKV replication compared to the media control; 56.8 μM ciprofloxacin added 2, 4, and 6 hours p.i. significantly reduced replication, and 116.1 μM ciprofloxacin added at 2, 4, 6, and 8 hours p.i. significantly reduced replication (full statistics in Table S1).

### Enoxacin treatment of ZIKV-infected mice did not alleviate or exacerbate weight loss

To evaluate the *in vivo* efficacy of enoxacin to suppress ZIKV, A129 mice were infected with ZIKV and treated with enoxacin in two independent experiments described in Fig. 1 and Table 2. In both experiments, all mice lost weight, irrespective of treatment (Fig. 3). Loss of weight by control mice was unexpected, and likely resulted from the effects of dosing these small (average 19.3 g) animals twice daily with 100 μL volume of liquid. In experiment 1, weight loss, quantified as the percent of initial weight, did not differ between ZIKV-infected mice treated with 10 mg/kg and 15 mg/kg (repeated measures ANOVA: F (5,55) = 0.7, P = 0.61) and mean percent weight lost by uninfected mice treated with 10 mg/kg and 15 mg/kg also did not differ significantly different from each other (repeated measures ANOVA: F (6,54) = 0.6, P = 0.70); therefore, we pooled the weight data by enoxacin treatment regardless of dosage for the ZIKV-infected and uninfected mice. There was a significant interaction between group (ZIKV-infected and enoxacin treated, ZIKV-infected and diluent treated, or Sham-infected and enoxacin treated) and day post infection (repeated measures ANOVA: F (12, 194) = 3.1, P = 0.0006). Pairwise comparisons with t-tests identified differences in weight loss on days 1, 3, 4, 5, and 6 p.i. (P < 0.05). On days 1, 3, 4, and 5 p.i. the percent of initial weight for the sham infected mice treated with enoxacin was lower compared to the ZIKV-infected mice treated with enoxacin (pairwise t-test P<0.05; for full statistics see Table S2). On day 6 p.i. the mean percent of initial weight of the uninfected mice treated with enoxacin was greater than that of the infected mice regardless of treatment (pairwise t-test P<0.05; for full statistics see Table S2). Additionally, at no point during the experiment did weight loss differ between the ZIKV infected mice treated with enoxacin and the diluent control mice (Fig. 3a, Table S2). In experiment 2, weight loss by day 6 was less drastic than, and significantly different from, experiment 1 (mean percent of initial weight (SE): experiment 1 = 84.0 (1.0), experiment 2 = 93.0 (0.9); t = −5.7, df = 55, P = 2.6×10^−7^), likely due to improved technical facility in dosing these very small mice, and weight loss was not significantly different among treatments (Fig. 3b, repeated measures ANOVA: F (18,132) = 1.4, P = 0.10).

**Fig 3.**
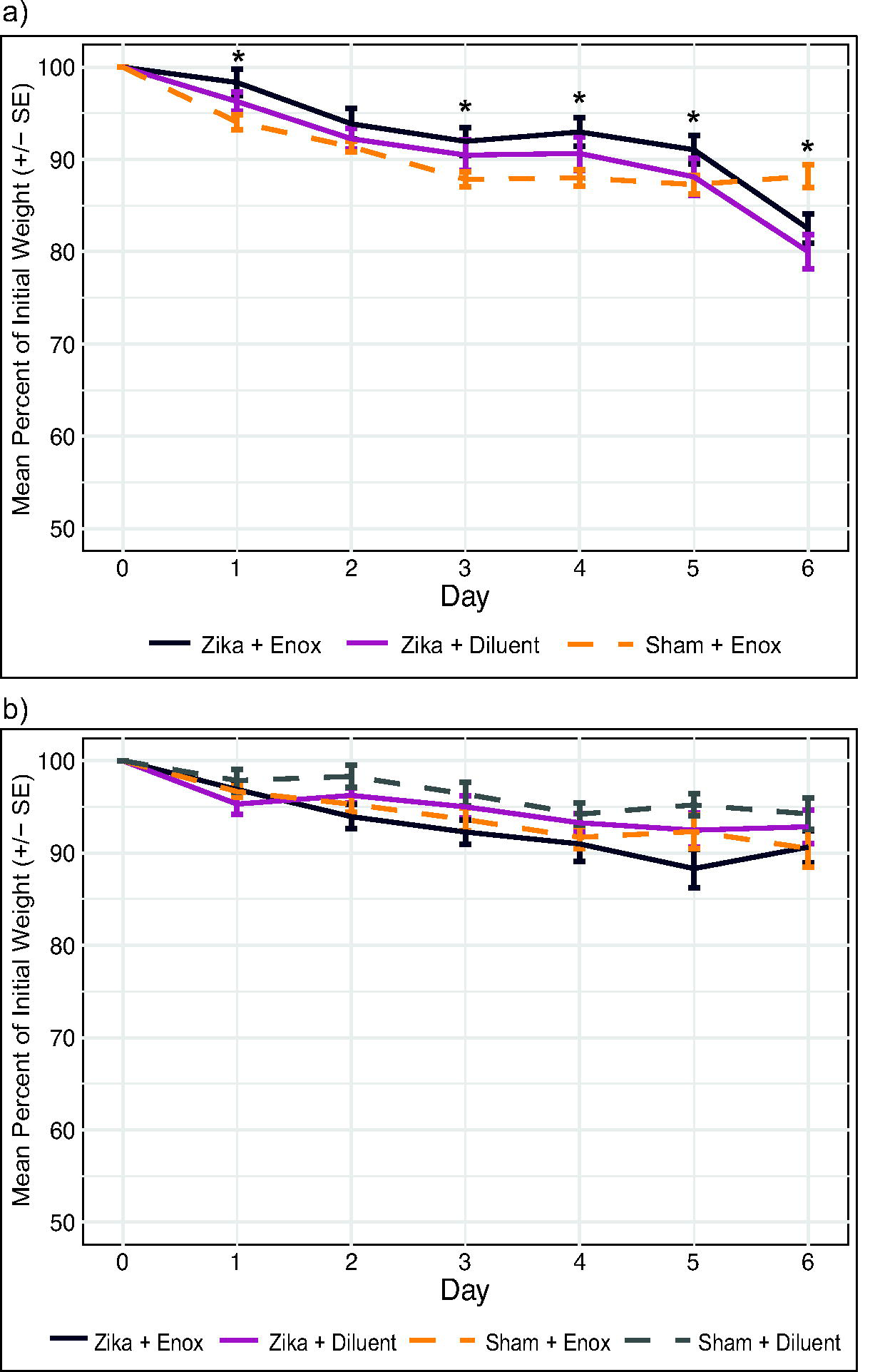
Mouse weight loss did not differ among treatments. (a) Daily percent of initial weight for experiment 1 was the same for ZIKV infected mice treated with enoxacin (10 mg/kg and 15 mg/kg combined) or diluent and uninfected mice treated with enoxacin (10 mg/kg and 15 mg/kg combined) until day 6 p.i. when the infected mice, regardless of treatment, lost significantly more weight than the uninfected controls. (b) Percent of initial weight for experiment 2 was not different among treatment groups. * at least one group is different at P <0.05. Sample sizes in Table 2; full statistics in Table S2.

### Enoxacin suppressed ZIKV replication in mouse testes, but not serum, brain, or liver

#### Experiment 1

In this experiment, mice were infected with 1×10^5^ pfu ZIKV and subsequently treated with enoxacin. ZIKV titer in the serum of mice treated with 15 mg/kg enoxacin was X-fold higher than those mice treated with 10 mg/kg enoxacin, a significant difference (Fig 4a; ANOVA F (2,17) = 4.7, P = 0.02; pairwise t-test P < 0.05). However, neither dose of enoxacin altered ZIKV titers in serum significantly relative to control mice (Fig 4a; pairwise t-test P = 0.22 for both). Similarly ZIKV titer in the brains of mice treated with 15 mg/kg enoxacin was approximately tenfold higher than that of mice treated with 10 mg/kg enoxacin (Fig. 4b; ANOVA F(2,17) = 4.2, P = 0.03; pairwise t-test P < 0.05), but these titers did not differ from the virus titer in brains of control mice (pairwise t-test, P = 0.73 for 10 mg/kg enoxacin and P = 0.06 for 15 mg/kg enoxacin). Given the small sample sizes of this study, it is possible that this effect is due to random sampling. In contrast to serum and brain, mean ZIKV titers in the testes of mice treated with 10 mg/kg and 15 mg/kg were not significantly different from each other (Fig. 4c; 5.4 log_10_ pfu/g (± 0.1 SE) vs 5.7 log_10_ pfu/g (± 0.1 SE); t = −1.6, df = 8, P = 0.14), and were both lower than the control group (6.2 log_10_ pfu/g (± 0.5 SE)), albeit only the decrease from 10mg/kg treatment was significant (ANOVA F(3,19) = 3.0, P = 0.05; pairwise t-test P < 0.05). To compensate for the decrease in sample size inherent in analyzing only males, the data from the two enoxacin concentrations were pooled. The mean ZIKV titer in testes for mice treated with any dosage of enoxacin was 5.5 log_10_ pfu/g (± 0.1 SE), which was significantly lower than 6.2 log_10_ pfu/g (± 0.5 SE) in the control group (pairwise t-test P < 0.05).

**Fig 4.**
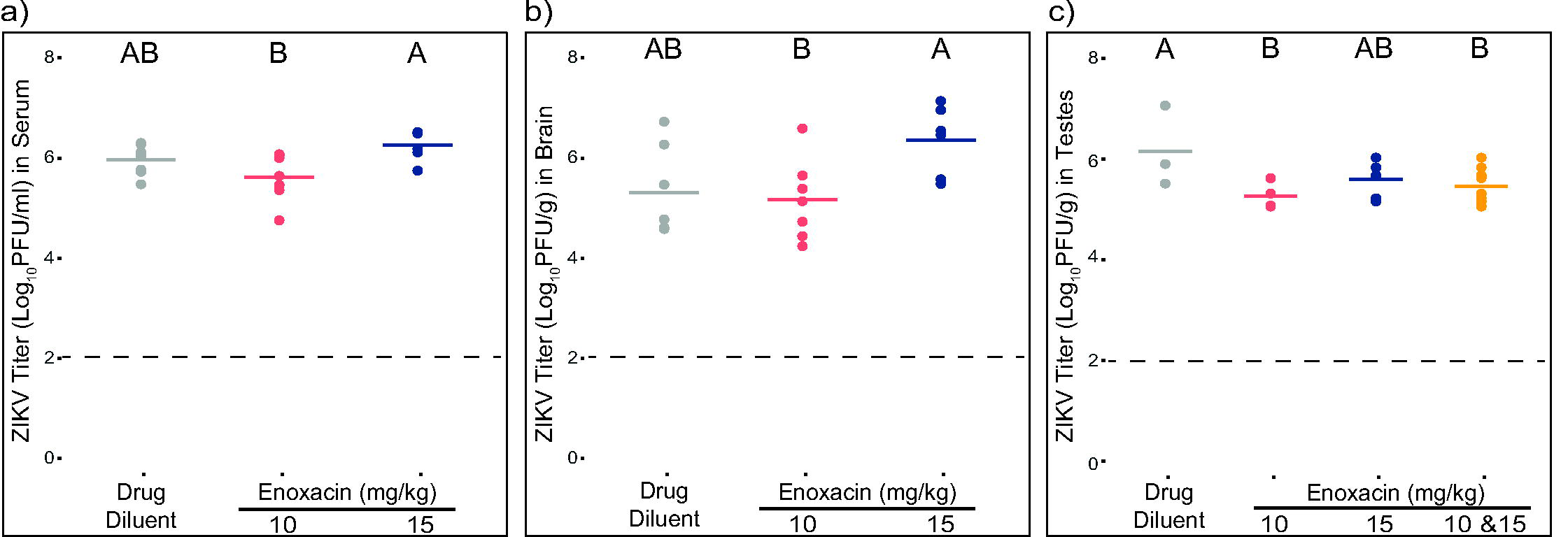
Treatment with enoxacin following high-titer infection suppresses ZIKV replication in mouse testes but not in sera or brain. Individual (dots) and mean (line) ZIKV titers of mice treated with the drug diluent or enoxacin from (a) sera, (b) brain, and (c) testes. Sample sizes for each treatment are listed in Table 2 and statistical analysis is described in the text.

#### Experiment 2

In experiment 2 mice were pre-treated with enoxacin after which they were infected with 1×10^2^ pfu ZIKV and subsequently dosed daily with enoxacin. As expected, ZIKV titers in serum and brain were two orders of magnitude lower than those in experiment 1, however titers in the testes were similar between the two experiments. Consistent with experiment 1, ZIKV titers in sera, brains, and livers of enoxacin-treated mice were not different from control mice (Fig. 5a-c, all P-values > 0.05), while ZIKV titer in testes of treated mice were 2-fold lower than those of control mice, and this difference was significant (Fig. 5d, pairwise t = −5.4, df = 5, P = 0.003).

**Fig 5.**
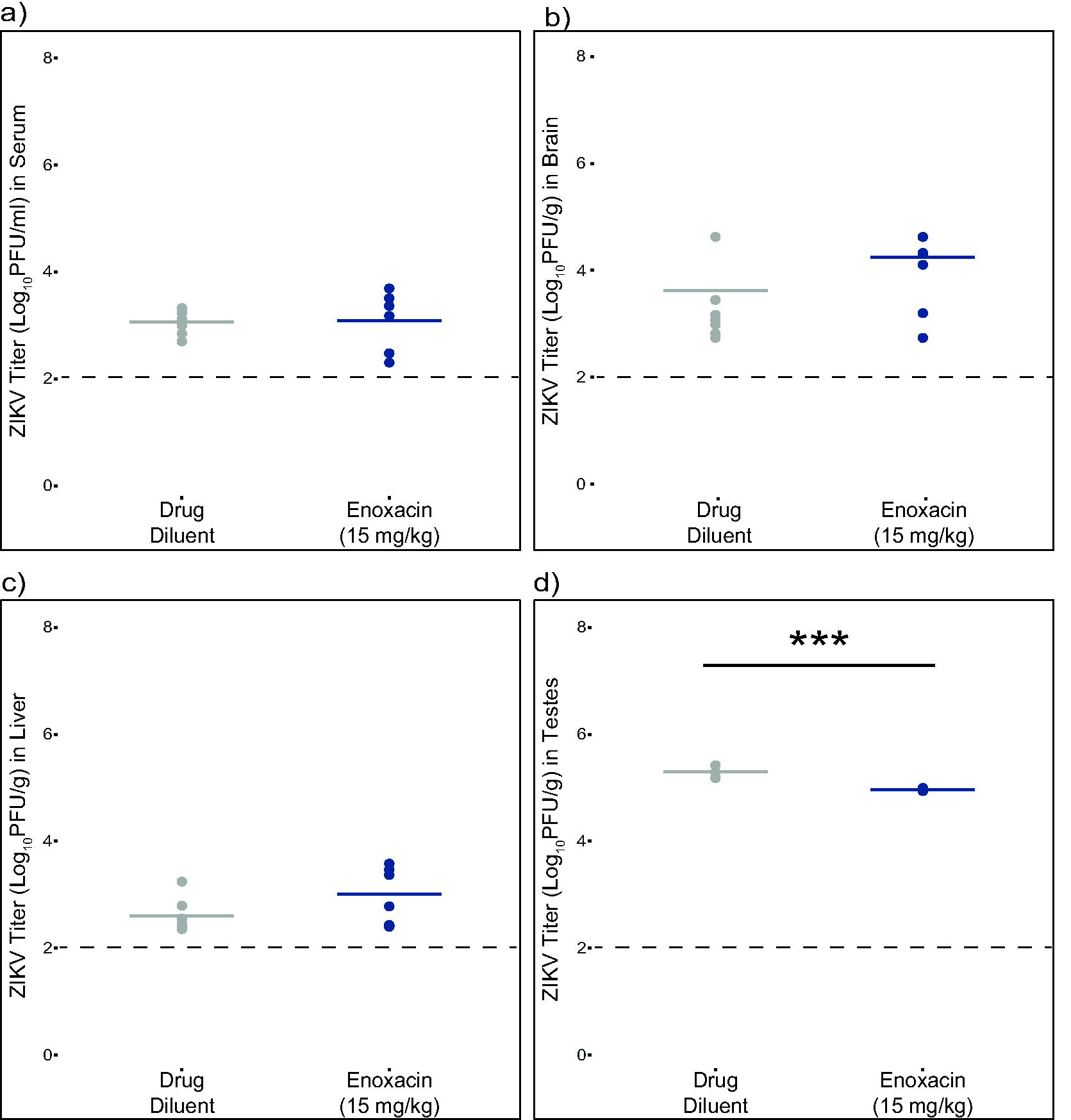
Treatment with enoxacin prior to and following low-titer ZIKV infection suppresses viral replication in the testes but not in serum, brain or liver. Individual (dots) and mean (line) ZIKV titers of mice treated with the drug diluent or enoxacin from (a) sera, (b) brain, (c) liver, and (d) testes. Sample sizes for each treatment are listed in Table 2 and statistical analysis is described in the text.

### Enoxacin does not inhibit ZIKV replication in mouse Sertoli cells

To investigate why the effect of enoxacin on ZIKV infection in mice was limited to the testes, the EC_50_ of this drug was quantified in both mouse Sertoli cells and HEK-293 cells at each of two MOI: 0.1 and 1.0. As expected, at 32 ° C, higher initial MOI generally resulted in higher ZIKV titers, particularly at early timepoints p.i., in both cell lines. Sertoli cells must be cultured at 32 ° C, so, for a fair comparison, the potency of enoxacin in HEK-293 cells was re-tested at 32 ° C. Moreover, to extend the window for comparison, virus was harvested at both 2 days (Fig. 6a) and at 5 days p.i. (Fig. 6b).

**Fig 6.**
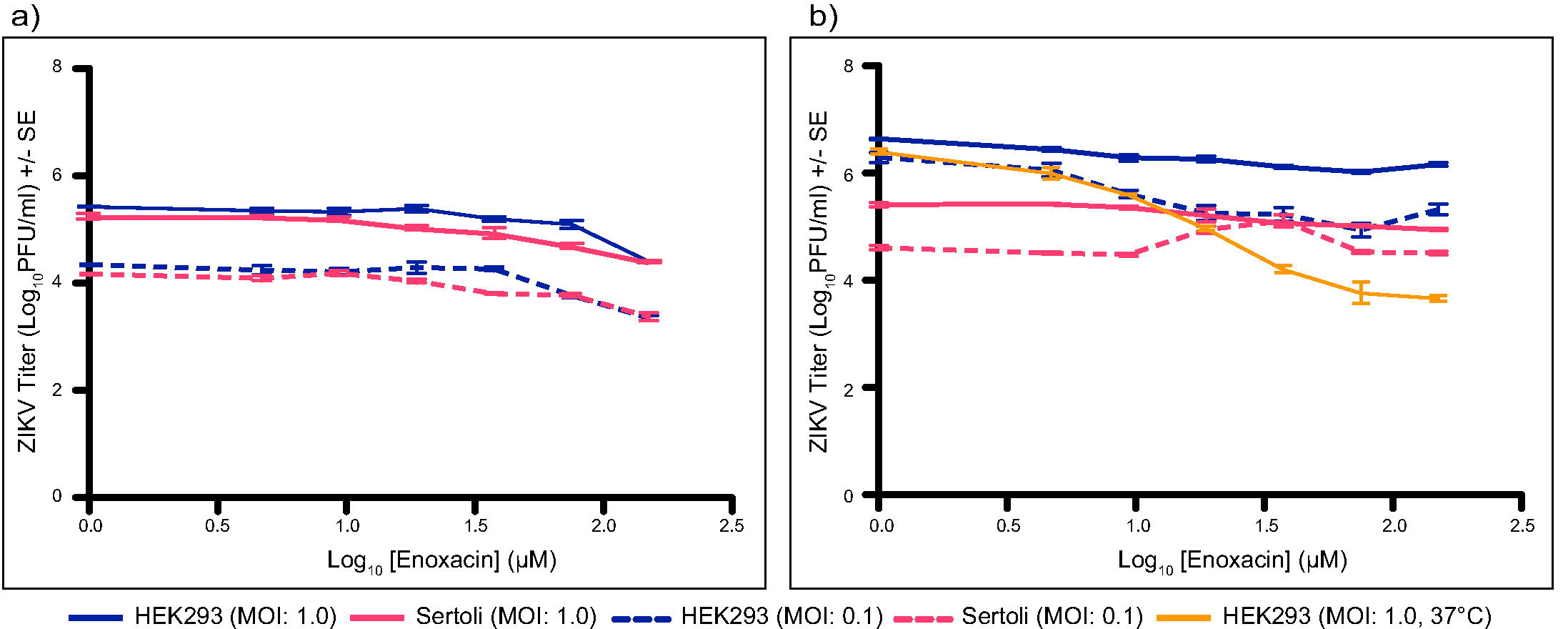
Enoxacin does not suppress ZIKV in mouse Sertoli cells two days pi (a) or five days pi (b). Dose-response curves for enoxacin and ZIKV titer at 32 ° C in Sertoli cells (pink) and HEK-293 cells (blue) at two MOI, 0.1 (dashed) and 1.0 (solid). Dose-response curve for enoxacin and ZIKV titer in HEK-293 cells at 37 ° C (yellow) is included for comparison on day 5 p.i. Statistical analysis is described in the text.

As expected, at 32 ° C, higher initial MOI generally resulted in higher ZIKV titers, particularly at early timepoints p.i., in both cell lines. Kumar *et al*., Siemann *et al*., and Mlera and Bloom have previously tested the replication of ZIKV in Sertoli cells and found them to be highly susceptible to ZIKV infection (59–61). In our study, in the absence of enoxacin, there was no difference in ZIKV replication in HEK-293 and Sertoli cells 2 days p.i. (linear model, β = −0.2, P = 0.59), but at 5 days p.i. ZIKV titers in Sertoli cells were significantly lower than in HEK-293 cells (linear model, β = −1.4, P = 7.4×10^−7^).

Unlike the dose response curve in HEK-293 cells incubated at 37 ° C, increasing concentration of enoxacin in both cell lines at 32 ° C did not result in a sharp inflection in ZIKV titer, making it difficult to accurately quantify the EC_50_ via non-linear regression. Instead, general linear models were used to test the relative potency in the two cell types at 32 ° C. Potency at 32 ° C in either cell line was detectable at the higher concentrations of enoxacin on day 2, but not on day 5 (Fig. 6a and 6b).

We tested two hypotheses for greater impact of the drug in testes: first that this effect may have been due to the lower temperature of the testes and second that it may have been due to a greater potency in testis cells. To test the first hypothesis, we compared enoxacin potency in HEK-293 cells infected with ZIKV at MOI 1 incubated at 32 ° C or 37 ° C and harvested on day 5 (data at 37 ° C was collected as part of the initial EC_50_ analysis). Counter to the hypothesis, enoxacin potency was greater at 37 ° C compared to 32 ° C in this cell line (linear model, β = −0.2, P = 4.7×10^−15^).

We tested the second hypothesis by comparing the impact of enoxacin in HEK-293 and Sertoli cells at 32 ° C. We used Sertoli cells as our model testis cell, while acknowledging that the testes are composed of many cell types and results from Sertoli cells cannot be generalized to the testes as a whole. In this analysis enoxacin concentration, cell type, and MOI and their interactions were all included in the model. On day 2 pi, interaction between enoxacin concentration and cell type was not significant (β = 0.01, P = 0.83), while on day 5 p.i. there was a significant interaction (β = 0.7, P = 0.0001). On both days, enoxacin concentration and Sertoli cells continue to negatively impact ZIKV replication (day 2 linear model, β = −0.4, P = 3.2×10^−12^; β = −0.2, P = 0.03, respectively; day 5 linear model, β = −0.8, P = 4.9×10^−11^; β = −1.4, P = 1.9×10^− 8^, respectively) meaning that ZIKV titer decreased as enoxacin concentration increased and ZIKV infection of Sertoli cells resulted in lower titers overall compared to HEK-293 cells. However, on day 5 the interaction between enoxacin concentration and Sertoli cells had a positive impact on ZIKV titer, meaning that enoxacin was less effective in Sertoli cells than in HEK-293 cells.

## Discussion

Flavivirus infections are acute, and treatment must be initiated rapidly to be effective (6,62,63). However, individuals infected with different flaviviruses often present with similar symptoms (62,64), and in many places where flavivirus infections are common, diagnostic capacity is limited (62,65). Thus, the ideal antiflaviviral drug will have broad efficacy across different members of the genus (12,66,67). We found that the three fluoroquinolones used in this study, enoxacin, ciprofloxacin and difloxacin, all suppressed replication of the six flaviviruses tested at low micromolar concentrations, with the exception that difloxacin lacked potency for MODV. These six flaviviruses, DENV-1, DENV-2, DENV-4, ZIKV, LGTV, and MODV, span the diversity of human pathogenic flaviviruses (68,69). While enoxacin consistently demonstrated the lowest EC_50_ values, difloxacin exhibited the best selectivity indexes. These findings suggest that fluoroquinolones could offer broad-spectrum anti-flaviviral activity, a very desirable property.

Though the anti-flaviviral mechanism of action of fluoroquinolones remains unknown, several possible mechanisms merit exploration. First, suppression of flaviviral replication by fluoroquinolones could be mediated by enhancement of RNAi (23,24). The current study was motivated by the discovery of this effect, and the drugs we chose to evaluate span the range from high (enoxacin) to low (difloxacin) impact on RNAi (23). Our finding that all three fluoroquinolones tested inhibited replication of the six flaviviruses tested suggests that the antiviral action of fluoroquinolones cannot be attributed solely to enhancement of RNAi.

Second, fluoroquinolones could prevent endocytosis-mediated viral entry. Fluoroquinolones are derived from the original quinolone, nalidixic acid, which is a biproduct of synthesizing chloroquine, an antimalarial drug (70). Consequently, fluoroquinolones and chloroquine share a 4-quinolone structure. As a weak base, chloroquine is known to inhibit viral entry by increasing the pH of vesicles required for endocytosis-mediated cellular entry (71–80). Chloroquine has been shown to suppress ZIKV and DENV in cultured mammalian cells, including Vero, HuH-7, U937, human neural progenitor (hNPC), and human brain microvascular endothelial cells (hBMEC) cells, with a range of EC_50_ values from 1 μM to 14 μM against ZIKV (79,81–84) which is quite similar to the range of ciprofloxacin EC_50_ values against ZIKV (6.5-13.4 µM) determined in this study. Initial studies of chloroquine in mouse and monkey models were promising (82,85,86), however, results from two human clinical trials with DENV found improvement in some dengue-associated symptoms, such as pain, but no reduction in viremia or infection duration (87,88). We speculate that the structural similarities of fluoroquinolones and chloroquine could be the shared basis of their anti-flaviviral efficacy (18). If chloroquine and fluoroquinolones share similar antiviral mechanisms of action, then results from studies of chloroquine could offer insight into what modifications could be made to the fluoroquinolones to increase their antiviral efficacy.

Third, fluoroquinolones could suppress the viral helicase as their mechanism of action. Khan et al. (32) demonstrated that many fluoroquinolones, including enoxacin and difloxacin, suppressed HCV replication and inhibited the viral helicase *in vitro* (32). Further studies will be needed to assess the generality and *in vivo* relevance of this result.

To distinguish among some of these potential mechanisms of action, we conducted a time-of-addition study of all three drugs. This approach has previously been used to reveal that 50 μM chloroquine, which is approximately 5 times the EC_50_ value, reduces viral RNA 64-fold when added at the time of infection, likely reflecting inhibition of viral entry. (79). In our study, we found that difloxacin and ciprofloxacin suppression activity was wider, encompassing 2 to 8 hours p.i. across assays, and in a subset of assays these drugs showed potency when cells were treated prior to infection, similar to chloroquine. In contrast, enoxacin suppression was restricted to 2 to 6 hours p.i. These results indicate that difloxacin and ciprofloxacin may impact early and intermediate viral life cycle stages whereas enoxacin’s effect is limited to intermediate stages. Combined, these data suggest that ciprofloxacin and difloxacin may share a mechanism of action that is different than that of enoxacin.

In light of enoxacin’s low EC_50_, we tested the ability of this drug mitigate weight loss (55) and suppress serum viremia and virus titer in key tissues including liver, brain and testes (55) in ZIKV-infected A129 mice via two experiments. First, we infected A129 mice with a high titer of ZIKV and then treated immediately after infection with drug diluent or 10 or 15 mg/kg of enoxacin, which for the average weight of a 5-week-old mouse corresponds to 6 or 10 μM enoxacin. These concentrations were selected because they are close to the EC_50_ value for enoxacin in HEK-293 cells (18.1 μM) and also within the range of peak human serum concentration after oral consumption of clinically available dosages of enoxacin, (5 to 11 μM) (56,57). Unexpectedly, all of the mice, including the sham-infected mice, lost weight in this experiment. Two non-exclusive explanations for the weight loss are that oral administration of 100 μL liquid twice a day caused satiety and prevented the mice from eating normally or that some component of the dose caused stomach discomfort which also prevented the mice from eating. Despite this, none of the mice lost more than 20% of their weight and therefore did not reach the cutoff for euthanasia. Enoxacin treatment had no impact on serum viremia or on virus titer in liver or brain. As fluoroquinolones readily cross the blood-brain barrier (89) and are metabolized in the liver (90), the absence of a drug effect in these tissues is unlikely to be due to lack of enoxacin availability. The most intriguing result of this experiment, however, was that male mice treated with enoxacin showed a significant 50% decrease in ZIKV titer in the testes.

To assess the reproducibility of these findings under a different treatment regimen, we next tested the effect of pre-treating mice with enoxacin and then infecting them with ZIKV. As in the first experiment, all mice lost weight (though weight loss was less than in the first experiment) and ZIKV titers in serum, liver and brain did not differ between enoxacin-treated and control mice. Importantly, consistent with the first experiment, ZIKV was significantly suppressed in the testes of enoxacin-treated mice relative to controls.

We initially hypothesized that the restriction of enoxacin’s efficacy to the testes *in vivo* was due to higher efficacy in specific cell types in the testes or the lower temperature of the testes. However, counter to these explanations, we found that enoxacin was less effective against ZIKV at 32°C than 37°C and less effective against ZIKV in Sertoli cells than in human kidney cells. However, Sertoli cells are one of many cell types in the testes, which also contain stem cells, spermatozoa and Leydig cells, which vary in their susceptibility to ZIKV (60,61,91– 93), thus our findings in Sertoli cells do not reveal the action of enoxacin in the testes as a whole. Immunohistochemical staining of ZIKV-infected mouse testes 7 days p.i. has revealed the presence of viral antigen primarily in the stem cells of the seminiferous tubules and in the seminal fluid from the vas deferens (93). Using *in situ* hybridization, viral RNA of a mouse-adapted strain of ZIKV was detected in the stem cells (spermatogonia and primary spermatocytes), Sertoli cells, and spermatozoa of ZIKV-infected mouse testes at 7 days p.i. (92). ZIKV-infected germ cells were detected in the basal layer of the seminiferous tubules of ZIKV-infected olive baboons via immunofluorescence 11 days p.i. (94).

Recently, Xu *et al*. (35) reported that ZIKV infection of hNPCs activates the RNAi antiviral response and elicits the production of virus-derived small interfering RNAs (vsi-RNA), but infection of human neurons does not, indicating that cellular differentiation degrades the functionality of RNAi. Additionally, Xu et al. (35) demonstrated that treatment with enoxacin, a known enhancer of RNAi (23,24), significantly suppressed ZIKV replication in hNPCs. The HEK-293 cells in which we demonstrated enoxacin efficacy against ZIKV have stem-cell like properties (95). Our time-of-addition assays indicate that enoxacin, ciprofloxacin, and difloxacin all suppress an intermediate life stage of ZIKV, which is consistent with enhancement of RNAi. Furthermore, we detected an impact of enoxacin in testes, which are rich in stem cells, but not in differentiated brain or liver cells. Thus, our results are generally consistent with those of Xu et al. (35), which implicate RNAi enhancement as an antiviral mechanism of enoxacin.

However, multiple alternative explanations for our *in vivo* findings must be considered. First and foremost, Xu et al. (35) used 10-fold more enoxacin (100 μM vs 10 μM) in their study than we used in ours. Additionally, testicular ZIKV infection results in oxidative stress, and antioxidants such as ebselen have been shown to reduce oxidative stress, lessen testicular damage, and prevent sexual transmission in mice (96). Like ebselen, fluoroquinolones are also known to act as antioxidants (27–29). Fluoroquinolones may also damage the testes and thereby restrict flavivirus replication. More research on the *in vivo* testicular toxicity of fluoroquinolones is needed, especially since ZIKV infection itself damages testicular tissues (92,97–99) although damage to human Sertoli cells is minimal (60).

More generally, several caveats pertain to our study. First, although use of the A129 immunocompromised mouse model to initially test compounds for *in vivo* efficacy against ZIKV has become a standard practice (82,100,101), nonetheless, these mice do lack an interferon response, and the interplay between interferon and ZIKV clearly shapes pathogenesis (102– 108). Thus, the reproducibility of our findings in other, immunocompetent animal models, such as the human STAT2 knockin mouse model or C57B1/6 mice treated with the anti-type I IFN receptor antibody (108–110), should be tested. Second, as we only tested enoxacin across two concentrations *in vivo*, a wider range of fluoroquinolones and fluoroquinolone concentrations should be investigated *in vivo* for efficacy in suppressing ZIKV. Third, our investigation of the testes-specific efficacy of enoxacin focused only on Sertoli cells, but efficacy in other testicular cells should also be evaluated, particularly stem cells.

In summary, we found that three fluoroquinolones had reasonable potency against six flaviviruses in cultured cells and enoxacin suppressed ZIKV titer in mouse testes. These results offer a foundation for further attempts to optimize fluoroquinolones to increase potency. Limiting replication of the virus in the testes is important, as ZIKV is capable of sexual transmission (111–114). The results from this study and that of Xu et al. (35) also suggest that testing the ability of fluoroquinolones to alleviate the teratogenic effects of ZIKV in relevant mouse models (109,115,116) should be a priority.

## Author Contributions

Conceptualization: SLPS CCA KAH SLR.

Investigation: SLPS CCA SRA EES EGJ KAH SLR.

Methodology: SLPS CCA KAH SLR.

Resources: RC JBA KAH SLR.

Supervision: KAH SLR.

Writing – original draft: SLPS KAH SLR.

Writing – review & editing: SLPS CCA RC SRA EES EGJ JBA KAH SLR.

## Supporting Information

**Table S1.** ANOVA results and pairwise t-test comparisons for ZIKV titer after treatment with enoxacin, difloxacin and ciprofloxacin from the time-of-addition assays in Fig. 2 and Fig. S3.

|  | Enoxacin |  | Difloxacin |  |  | Ciprofloxacin |  |  |
| --- | --- | --- | --- | --- | --- | --- | --- | --- |
| Concentration (μM) | 18.1 | 24.4 | 35.9 | 25.4 | 50.0 | 116.1 | 56.8 | 25.0 |
| MOI | 1.0 | 0.2 | 0.2 | 1.0 | 1.0 | 0.2 | 1.0 | 1.0 |
| F | 2.6 | 3.6 | 2.8 | 1.0 | 12.8 | 14.7 | 41.2 | 16.2 |
| DF | 8, 18 | 8, 18 | 8, 18 | 8, 18 | 8, 18 | 8, 18 | 8, 18 | 8, 18 |
| P | 0.04 | 0.01 | 0.04 | 0.45 | 5.2e-6 | 1.9e-6 | 5.2e-10 | 2.3e-7 |
| Time-of-addition (hour p.i.) |  |  |  |  |  |  |  |  |
| Media | AB | AB | A | A | A | A | AB | A |
| -2 | AB | AB | AB | A | BC | A | B | CD |
| 0 | A | ABC | AB | A | CD | A | B | BC |
| 2 | BC | CD | B | A | CD | B | D | E |
| 4 | BC | D | AB | A | D | B | CD | E |
| 6 | C | CD | B | A | D | B | C | DE |
| 8 | ABC | BCD | B | A | CD | B | B | DE |
| 12 | ABC | AB | B | A | A | A | A | BC |
| 18 | A | A | B | A | AB | A | AB | B |

**Table S2.** Pairwise t-test comparisons of mean percent weight change from *in vivo* ZIKV infection in Fig 3a

| Virus + Treatment | Day 0 | Day 1 | Day 2 | Day 3 | Day 4 | Day 5 | Day 6 |
| --- | --- | --- | --- | --- | --- | --- | --- |
| Sham + enoxacin | JK | FHI | DEG | BC | BC | B | BC |
| Zika + lactic acid | K | IJ | EFGH | BCDEFGH | BCDEFGH | BCD | A |
| Zika + enoxacin | JK | JK | GHI | DEFGH | EFGHI | CDEF | A |

**Figure S1.**
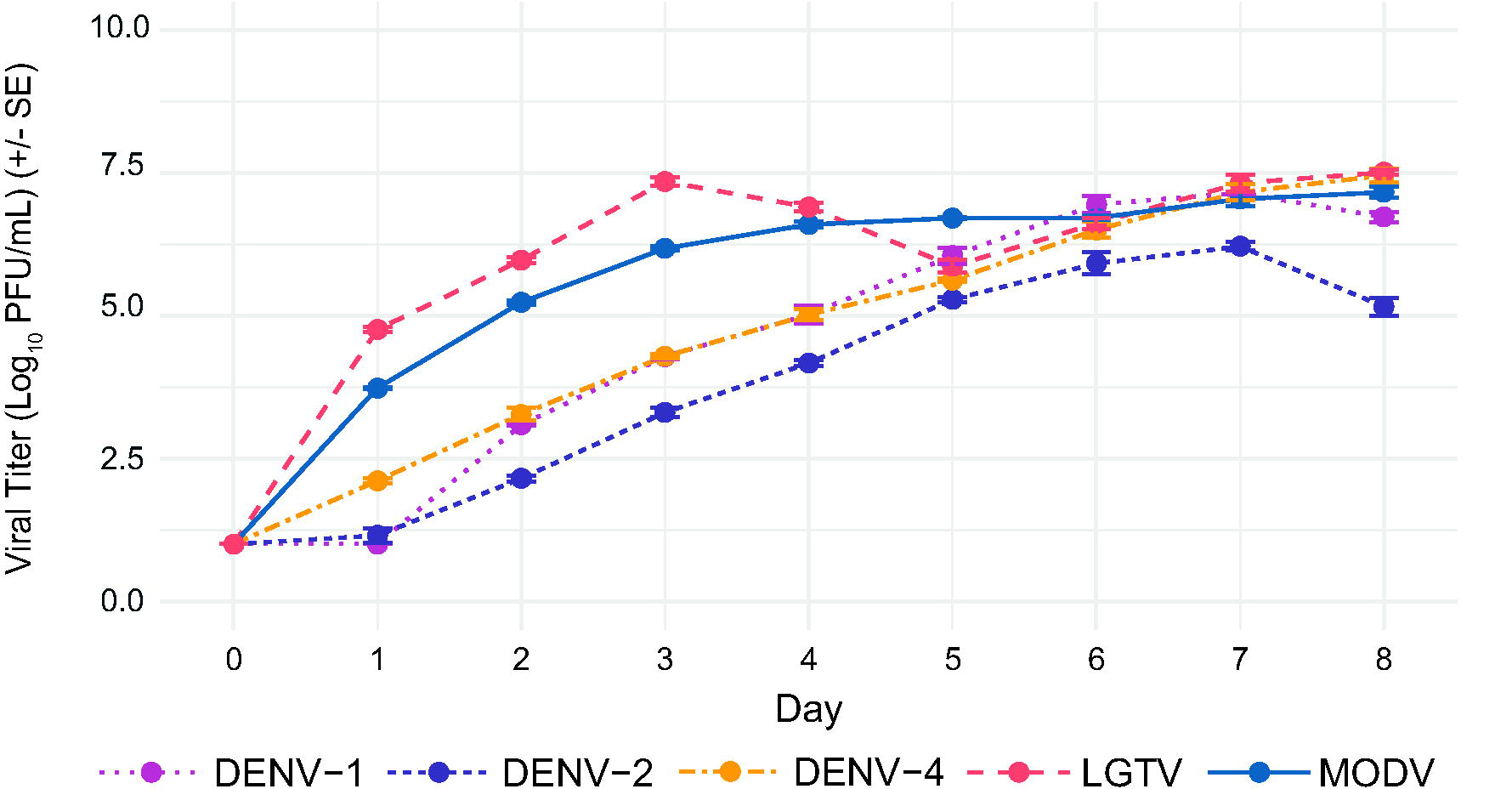
Replication kinetics of DENV-1, DENV-2, DENV-4, LGTV, and MODV at MOI 0.05 in HEK-293 cells.

**Figure S2.**
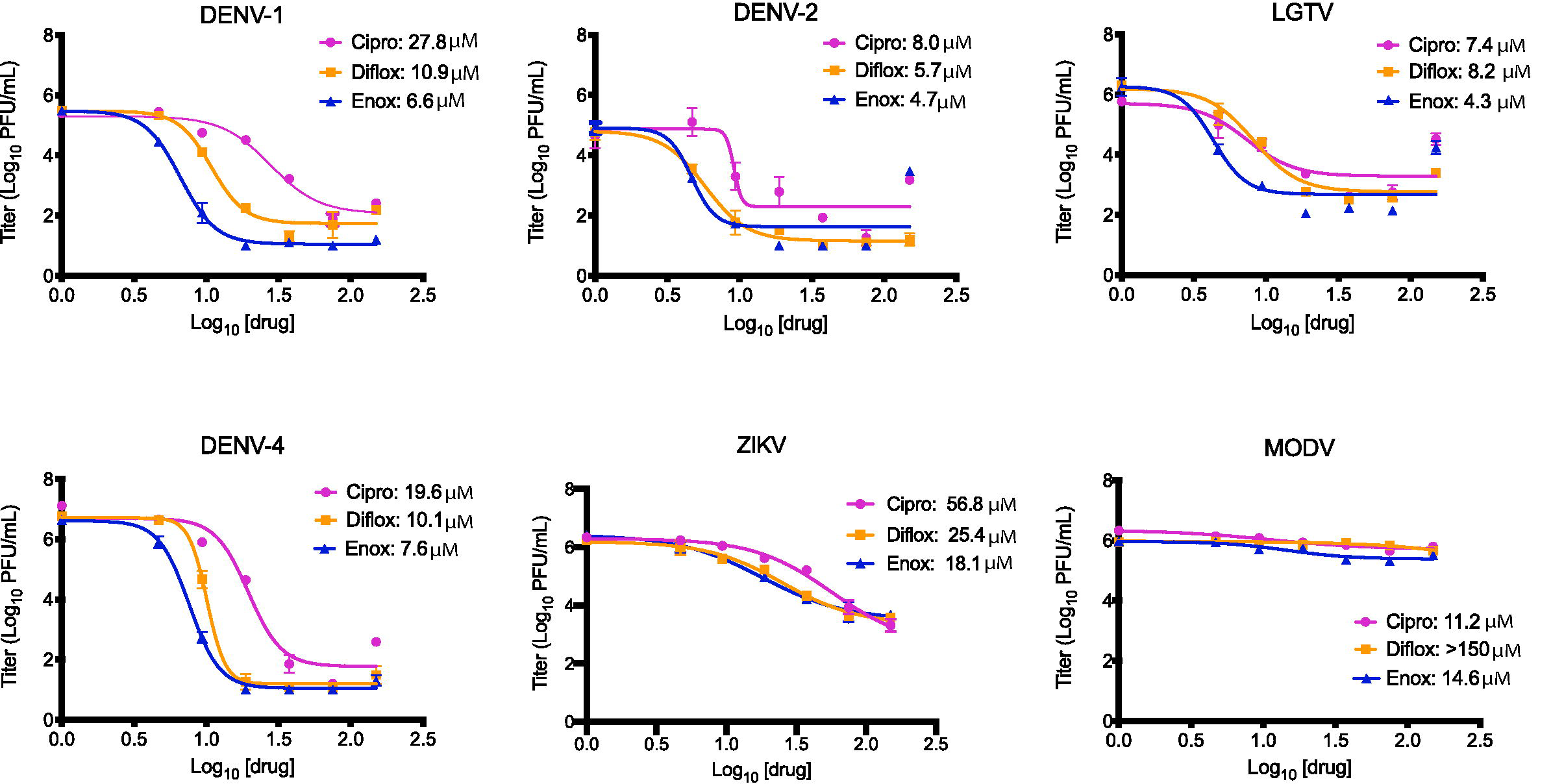
Dose response curves and EC_50_ for ciprofloxacin (pink), difloxacin (yellow), and enoxacin (blue) inhibition of DENV-1, DENV-2, DENV-4, ZIKV, LGTV, and MODV.

**Figure S3.**
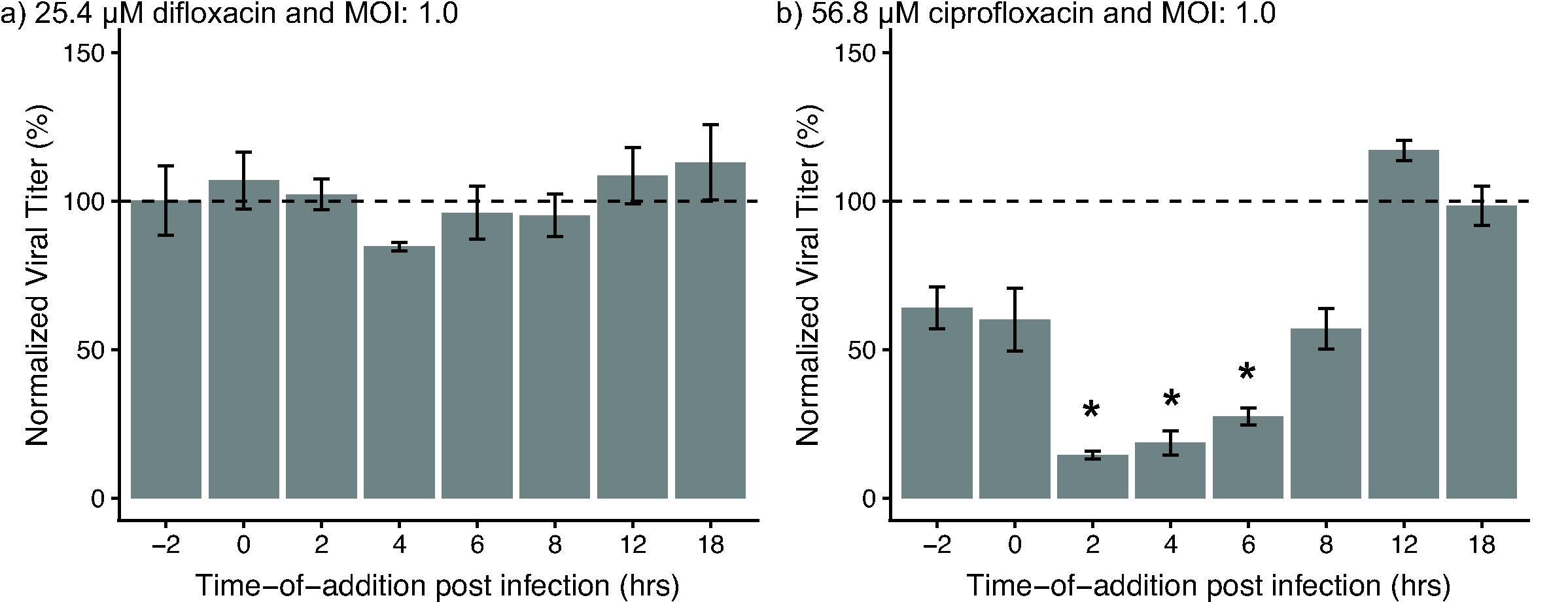
Impact on ZIKV replication by 25.4 μM difloxacin (a) and 56.8 μM ciprofloxacin (b) when added at designated timepoints. Viral titers (n = 3 replicates per drug per time point) for each time point were normalized to the average viral titer with media treatment and reported as average percent (titer at time point/average media titer*100). Differences in mean viral titers (log_10_ pfu/mL) were detected with ANOVA and pairwise t-tests; full pairwise statistics in Table S1. * P < 0.05 compared to media control.

**Data S1**. Excel spreadsheet containing the underlying data for Figures 2-6, Figures S1-S3, Table 3, Table S1 and S2.

## References

1. Bhatt S, Gething PW, Brady OJ, Messina JP, Farlow AW, Moyes CL, et al. The global distribution and burden of dengue. Nature. 2013;496:504–7.

2. Mlakar J, Korva M, Tul N, Popović M, Poljšak-Prijatelj M, Mraz J, et al. Zika Virus Associated with Microcephaly. N Engl J Med. 2016;

3. Honein MA, Dawson AL, Petersen EE, Jones AM, Lee EH, Yazdy MM, et al. Birth Defects Among Fetuses and Infants of US Women With Evidence of Possible Zika Virus Infection During Pregnancy. JAMA. 2016;30333(1):59–68.

4. Cao-Lormeau V-M, Blake A, Mons S, Lastère S, Roche C, Vanhomwegen J, et al. Guillain-Barré Syndrome outbreak associated with Zika virus infection in French Polynesia: a case-control study. Lancet. 2016;387(10027):1531–9.

5. Ružek D, Dobler G, Mantke OD. Tick-borne encephalitis: pathogenesis and clinical implications. Travel Med Infect Dis. 2010;8(4):223–32.

6. Kemenesi G, Bányai K. Tick-borne flaviviruses, with a focus on Powassan virus. Clin Microbiol Rev. 2018;32(1):e00106–17.

7. Weaver SC, Costa F, Garcia-Blanco MA, Ko AI, Ribeiro GS, Saade G, et al. Zika virus: History, emergence, biology, and prospects for control. Antiviral Res. 2016;130:69–80.

8. Le Flohic G, Porphyre V, Barbazan P, Gonzalez J-P. Review of climate, landscape, and viral genetics as drivers of the Japanese encephalitis virus ecology. PLoS Negl Trop Dis. 2013;7(9):e2208.

9. Petersen LR, Brault AC, Nasci RS. West Nile virus: review of the literature. JAMA. 2013;310(3):308–15.

10. Mackenzie JS, Gubler DJ, Petersen LR. Emerging flaviviruses: the spread and resurgence of Japanese encephalitis, West Nile and dengue viruses. Nat Med. 2004;10(12s):S98.

11. Ishikawa T, Yamanaka A, Konishi E. A review of successful flavivirus vaccines and the problems with those flaviviruses for which vaccines are not yet available. Vaccine. 2014;32(12):1326–37.

12. Boldescu V, Behnam MAM, Vasilakis N, Klein CD. Broad-spectrum agents for flaviviral infections: dengue, Zika and beyond. Nat Rev Drug Discov. 2017;16(8):565.

13. Saiz J-C. Therapeutic Advances Against ZIKV: A Quick Response, a Long Way to Go. Pharmaceuticals. 2019;12(3):127.

14. Lim S, Shi P-Y. West Nile virus drug discovery. Viruses. 2013;5(12):2977–3006.

15. Lim SP, Wang Q-Y, Noble CG, Chen Y-L, Dong H, Zou B, et al. Ten years of dengue drug discovery: progress and prospects. Antiviral Res. 2013;100(2):500–19.

16. Kaptein SJF, Neyts J. Towards antiviral therapies for treating dengue virus infections. Curr Opin Pharmacol. 2016;30:1–7.

17. Zou J, Shi P-Y. Strategies for Zika drug discovery. Curr Opin Virol. 2019;35:19–26.

18. Wolfson JS, Hooper DC. The Fluoroquinolones: Structures, Mechanisms of Action and Resistance, and Spectra of Activity In Vitro. Antimicrob Agents Chemother. 1985;28(4):581–6.

19. Kohanski MA, Dwyer DJ, Collins JJ. How antibiotics kill bacteria: from targets to networks. Nat Rev Microbiol. 2010;8(6):423.

20. Lindenbach BD, Rice CM. Molecular biology of flaviviruses. Adv Virus Res. 2003;59:23–62.

21. Nagy PD, Pogany J. The dependence of viral RNA replication on co-opted host factors. Nat Rev Microbiol. 2012;10(2):137.

22. Melo S, Villanueva A, Moutinho C, Davalos V, Spizzo R, Ivan C, et al. Small molecule enoxacin is a cancer-specific growth inhibitor that acts by enhancing TAR RNA-binding protein 2-mediated microRNA processing. Proc Natl Acad Sci. 2011;108(11):4394–9.

23. Shan G, Li Y, Zhang J, Li W, Szulwach KE, Duan R, et al. A small molecule enhances RNA interference and promotes microRNA processing. Nat Biotechnol. 2008;26(8):933–40.

24. Zhang Q, Zhang C, Xi Z. Enhancement of RNAi by a small molecule antibiotic enoxacin. Cell Res. 2008;18(10):1077.

25. Sharma BN, Li R, Bernhoff E, Gutteberg TJ, Rinaldo CH. Fluoroquinolones inhibit human polyomavirus BK (BKV) replication in primary human kidney cells. Antiviral Res. 2011;92(1):115–23.

26. Simon N, Bochman ML, Seguin S, Brodsky JL, Seibel WL, Schwacha A. Ciprofloxacin is an inhibitor of the Mcm2-7 replicative helicase. Biosci Rep. 2013;33(5):e00072.

27. Dalhoff A. Immunomodulatory activities of fluoroquinolones. Infection. 2005;33(2):55–70.

28. Tazi KA, Moreau R, Hervé P, Dauvergne A, Cazals-Hatem D, Bert F, et al. Norfloxacin reduces aortic NO synthases and proinflammatory cytokine up-regulation in cirrhotic rats: role of Akt signaling. Gastroenterology. 2005;129(1):303–14.

29. Akamatsu H, Niwa Y, Sasaki H, Matoba Y, Asada Y, Horio T. Effect of pyridone carboxylic acid anti-microbials on the generation of reactive oxygen species in vitro. J Int Med Res. 1996;24(4):345–51.

30. Poon IKH, Chiu Y-H, Armstrong AJ, Kinchen JM, Juncadella IJ, Bayliss DA, et al. Unexpected link between an antibiotic, pannexin channels and apoptosis. Nature. 2014;507(7492):329.

31. Fukumoto R, Cary LH, Gorbunov N V, Lombardini ED, Elliott TB, Kiang JG. Ciprofloxacin modulates cytokine/chemokine profile in serum, improves bone marrow repopulation, and limits apoptosis and autophagy in ileum after whole body ionizing irradiation combined with skin-wound trauma. PLoS One. 2013;8(3):e58389.

32. Khan IA, Siddiqui S, Rehmani S, Kazmi SU, Ali SH. Fluoroquinolones inhibit HCV by targeting its helicase. Antivir Ther. 2012;17:467–76.

33. Kojima H, Kaita KDE, Hawkins K, Uhanova J, Minuk GY. Use of fluoroquinolones in patients with chronic hepatitis C virus-induced liver failure. Antimicrob Agents Chemother. 2002;46(10):3280–2.

34. Yamaya M, Nishimura H, Hatachi Y, Yasuda H, Deng X, Sasaki T, et al. Levofloxacin inhibits rhinovirus infection in primary cultures of human tracheal epithelial cells. Antimicrob Agents Chemother. 2012;56(8):4052–61.

35. Xu YP, Qiu Y, Zhang B, Chen G, Chen Q, Wang M, et al. Zika virus infection induces RNAi-mediated antiviral immunity in human neural progenitors and brain organoids. Cell Res. 2019;29(4):265–73.

36. Durbin AP, Karron RA, Sun W, Vaughn DW, Reynolds MJ, Perreault JR, et al. Attenuation and immunogenicity in humans of a live dengue virus type-4 vaccine candidate with a 30 nucleotide deletion in its 3′-untranslated region. Am J Trop Med Hyg. 2001;65(5):405–13.

37. Hanley KA, Nelson JT, Schirtzinger EE, Whitehead SS, Hanson CT. Superior infectivity for mosquito vectors contributes to competitive displacement among strains of dengue virus. BCM Ecol. 2008;8(1).

38. Pletnev AG. Infectious cDNA clone of attenuated Langat tick-borne flavivirus (strain E5) and a 3′ deletion mutant constructed from it exhibit decreased neuroinvasiveness in immunodeficient mice. Virology. 2001;282(2):288–300.

39. Smith DR, Sprague TR, Hollidge BS, Valdez SM, Padilla SL, Bellanca SA, et al. African and Asian Zika virus isolates display phenotypic differences both in vitro and in vivo. Am J Trop Med Hyg. 2018;98(2):432–44.

40. Yin Z, Chen Y-L, Schul W, Wang Q-Y, Gu F, Duraiswamy J, et al. An adenosine nucleoside inhibitor of dengue virus. Proc Natl Acad Sci. 2009;106(48):20435–9.

41. Srivarangkul P, Yuttithamnon W, Suroengrit A, Pankaew S, Hengphasatporn K, Rungrotmongkol T, et al. A novel flavanone derivative inhibits dengue virus fusion and infectivity. Antiviral Res. 2018;151:27–38.

42. Desmyter J, Melnick JL, Rawls WE. Defectiveness of interferon production and of rubella virus interference in a line of African green monkey kidney cells (Vero). J Virol. 1968;2(10):955–61.

43. Emeny JM, Morgan MJ. Regulation of the interferon system: evidence that Vero cells have a genetic defect in interferon production. J Gen Virol. 1979;43(1):247–52.

44. Tarantino D, Cannalire R, Mastrangelo E, Croci R, Querat G, Barreca ML, et al. Targeting flavivirus RNA dependent RNA polymerase through a pyridobenzothiazole inhibitor. Antiviral Res. 2016;134:226–35.

45. Chuang F-K, Liao C-L, Hu M-K, Chiu Y-L, Lee A-R, Huang S-M, et al. Antiviral Activity of Compound L3 against Dengue and Zika Viruses In Vitro and In Vivo. Int J Mol Sci. 2020;21(11):4050.

46. Qing M, Zou G, Wang QY, Xu HY, Dong H, Yuan Z, et al. Characterization of dengue virus resistance to brequinar in cell culture. Antimicrob Agents Chemother. 2010;54(9):3686–95.

47. Xie X, Wang Q-Y, Xu HY, Qing M, Kramer L, Yuan Z, et al. Inhibition of dengue virus by targeting viral NS4B protein. J Virol. 2011;85(21):11183–95.

48. Byrd CM, Grosenbach DW, Berhanu A, Dai D, Jones KF, Cardwell KB, et al. Novel benzoxazole inhibitor of dengue virus replication that targets the NS3 helicase. Antimicrob Agents Chemother. 2013;57(4):1902–12.

49. Byrd CM, Dai D, Grosenbach DW, Berhanu A, Jones KF, Cardwell KB, et al. A novel inhibitor of dengue virus replication that targets the capsid protein. Antimicrob Agents Chemother. 2013;57(1):15–25.

50. Kato F, Ishida Y, Oishi S, Fujii N, Watanabe S, Vasudevan SG, et al. Novel antiviral activity of bromocriptine against dengue virus replication. Antiviral Res. 2016;131:141–7.

51. Pierson TC, Diamond M. Flaviviruses. In: Fields Virology: Sixth Edition. Wolters Kluwer Health Adis (ESP); 2013.

52. Bedos J-P, Azoulay-Dupuis E, Moine P, Muffat-Joly M, Veber B, Pocidalo J-J, et al. Pharmacodynamic activities of ciprofloxacin and sparfloxacin in a murine pneumococcal pneumonia model: relevance for drug efficacy. J Pharmacol Exp Ther. 1998;286(1):29–35.

53. Chartrand SA, Scribner RK, Marks MI, Dice J. Enoxacin pharmacokinetics and efficacy in CF-1 mice. J Antimicrob Chemother. 1987;19:221–4.

54. Abd El-Aty AM, Goudah A, Ismail M, Shimoda M. Disposition kinetics of difloxacin in rabbit after intravenous and intramuscular injection of Dicural. Vet Res Commun. 2005;29:297–304.

55. Rossi SL, Tesh RB, Azar SR, Muruato AE, Hanley KA, Auguste AJ, et al. Characterization of a novel murine model to study zika virus. Am J Trop Med Hyg. 2016;

56. Chang T, Black A, Dunky A, Wolf R, Sedman A, Latts J, et al. Pharmacokinetics of intravenous and oral enoxacin in healthy volunteers. J Antimicrob Chemother. 1988;21(suppl B):49–56.

57. Naber KG, Bartosik-Wich B, Sörgel F, Gutzler F. In vitro activity, pharmacokinetics, clinical safety and therapeutic efficacy of enoxacin in the treatment of patients with complicated urinary tract infections. Infection. 1985;13(5):219–24.

58. RCoreTeam. R: A language and environment for statistical computing. R foundation for statistical computing, Vienna, Austria. 2016.

59. Kumar A, Jovel J, Lopez-Orozco J, Limonta D, Airo AM, Hou S, et al. Human sertoli cells support high levels of zika virus replication and persistence. Sci Rep. 2018;8(1):1–11.

60. Siemann DN, Strange DP, Maharaj PN, Shi P-Y, Verma S. Zika virus infects human Sertoli cells and modulates the integrity of the in vitro blood-testis barrier model. J Virol. 2017;91(22).

61. Mlera L, Bloom ME. Differential Zika Virus Infection of Testicular Cell Lines. Viruses. 2019;11(1):42.

62. Katzelnick LC, Coloma J, Harris E. Dengue: knowledge gaps, unmet needs, and research priorities. Lancet Infect Dis. 2017;17(3):e88–100.

63. WHO. Dengue: guidelines for diagnosis, treatment, prevention and control. World Health Organization; 2009.

64. Sips GJ, Wilschut J, Smit JM. Neuroinvasive flavivirus infections. Rev Med Virol. 2012;22(2):69–87.

65. Racsa LD, Kraft CS, Olinger GG, Hensley LE. Viral hemorrhagic fever diagnostics. Clin Infect Dis. 2015;62(2):214–9.

66. Debing Y, Neyts J, Delang L. The future of antivirals: Broad-spectrum inhibitors. Curr Opin Infect Dis. 2015;28(6):596–602.

67. Vigant F, Santose MC, Lee N. Broad-spectrum antivirals against viral fusion. Nat Rev Microbiol. 2015;13(7):426–37.

68. Blitvich BJ, Firth AE. Insect-specific flaviviruses: a systematic review of their discovery, host range, mode of transmission, superinfection exclusion potential and genomic organization. Viruses. 2015;7(4):1927–59.

69. de Bernardi Schneider A, Machado DJ, Janies DA. Enhanced genome annotation strategy provides novel insights on the phylogeny of Flaviviridae. bioRxiv. 2019;674333.

70. Emmerson AM, Jones AM. The quinolones: decades of development and use. J Antimicrob Chemother. 2003;51(90001):13–20.

71. Helenius A, Marsh M, White J. Inhibition of Semliki Forest virus penetration by lysosomotropic weak bases. J Gen Virol. 1982;58(1):47–61.

72. Colpitts TM, Moore AC, Kolokoltsov AA, Davey RA. Venezuelan equine encephalitis virus infection of mosquito cells requires acidification as well as mosquito homologs of the endocytic proteins Rab5 and Rab7. Virology. 2007;369(1):78–91.

73. Zeichardt H, Wetz K, Willingmann P, Habermehl KO. Entry of poliovirus type 1 and Mouse Elberfeld (ME) virus into HEp-2 cells: Receptor-mediated endocytosis and endosomal or lysosomal uncoating. J Gen Virol. 1985;66(3):483–92.

74. Kronenberger P, Vrijsen R, Boeye A. Chloroquine induces empty capsid formation during poliovirus eclipse. JVirol. 1991;65(12):7008–11.

75. Khan M, Santhosh SR, Tiwari M, Lakshmana Roa PV, Parida M. Assessment of in vitro prophylactic and therapeutic efficacy of chloroquine against chikungunya virus in Vero cells. J Med Virol. 2010;88:817–24.

76. Nuckols JT, McAuley AJ, Huang YJS, Horne KM, Higgs S, Davey RA, et al. pH-Dependent entry of chikungunya virus fusion into mosquito cells. Virol J. 2014;11(1):4–7.

77. De Duve C, De Barsy T, Poole B, Tulkens P. Lysosomotropic agents. Biochem Pharmacol. 1974;23(18):2495–531.

78. Savarino A, Boelaert JR, Cassone A, Majori G, Cauda R. Effects of chloroquine on viral infections: an old drug against today’s diseases. Lancet Infect Dis. 2003;3(11):722–7.

79. Delvecchio R, Higa LM, Pezzuto P, Valadão AL, Garcez PP, Monteiro FL, et al. Chloroquine, an endocytosis blocking agent, inhibits zika virus infection in different cell models. Viruses. 2016;8(12):1–15.

80. Browning DJ. Pharmacology of chloroquine and hydroxychloroquine. In: Hydroxychloroquine and Chloroquine Retinopathy. Springer; 2014. p. 35–63.

81. Shiryaev SA, Mesci P, Pinto A, Fernandes I, Sheets N, Shresta S, et al. Repurposing of the anti-malaria drug chloroquine for Zika Virus treatment and prophylaxis. Sci Rep. 2017;7(1):1–9.

82. Li C, Zhu X, Ji X, Quanquin N, Deng YQ, Tian M, et al. Chloroquine, a FDA-approved Drug, Prevents Zika Virus Infection and its Associated Congenital Microcephaly in Mice. EBioMedicine. 2017;24:189–94.

83. Farias KJS, Machado PRL, da Fonseca BAL. Chloroquine inhibits dengue virus type 2 replication in Vero cells but not in C6/36 cells. Sci World J. 2013;2013.

84. Farias KJS, Machado PRL, de Almeida Junior RF, de Aquino AA, da Fonseca BAL. Chloroquine interferes with dengueL2 virus replication in U937 cells. Microbiol Immunol. 2014;58(6):318–26.

85. Cao B, Parnell LA, Diamond MS, Mysorekar IU. Inhibition of autophagy limits vertical transmission of Zika virus in pregnant mice. J Exp Med. 2017;214(8):2303–13.

86. Farias KJS, Machado PRL, Muniz JAPC, Imbeloni AA, da Fonseca BAL. Antiviral Activity of Chloroquine Against Dengue Virus Type 2 Replication in Aotus Monkeys. Viral Immunol. 2015;28(3):161–9.

87. Tricou V, Minh NN, Van TP, Lee SJ, Farrar J, Wills B, et al. A randomized controlled trial of chloroquine for the treatment of dengue in Vietnamese adults. PLoS Negl Trop Dis. 2010;4(8):e785.

88. Borges MC, Castro LA, Fonseca BAL da. Chloroquine use improves dengue-related symptoms. Mem Inst Oswaldo Cruz. 2013;108(5):596–9.

89. Scheld WM. Quinolone therapy for infections of the central nervous system. Rev Infect Dis. 1989;11(Suppl 5):S1194–202.

90. Outman WR. Metabolism and the fluoroquinolones. Am J Med. 1989;87(6C):37S–42S.

91. McDonald EM, Duggal NK, Brault AC. Pathogenesis and sexual transmission of Spondweni and Zika viruses. PLoS Negl Trop Dis. 2017;11(10):1–13.

92. Govero J, Esakky P, Scheaffer SM, Fernandez E, Drury A, Platt DJ, et al. Zika virus infection damages the testes in mice. Nature. 2016;540(7633):438.

93. Kawiecki AB, Mayton EH, Dutuze MF, Goupil BA, Langohr IM, Del Piero F, et al. Tissue tropisms, infection kinetics, histologic lesions, and antibody response of the MR766 strain of Zika virus in a murine model. Virol J. 2017;14(1):1–10.

94. Peregrine J, Gurung S, Lindgren MC, Husain S, Zavy MT, Myers DA, et al. Zika virus infection, reproductive organ targeting, and semen transmission in the male olive baboon. J Virol. 2019;94(1).

95. Debeb BG, Zhang X, Krishnamurthy S, Gao H, Cohen E, Li L, et al. Characterizing cancer cells with cancer stem cell-like features in 293T human embryonic kidney cells. Mol Cancer. 2010;9(1):180.

96. Simanjuntak Y, Liang JJ, Chen SY, Li JK, Lee YL, Wu HC, et al. Ebselen alleviates testicular pathology in mice with Zika virus infection and prevents its sexual transmission. PLoS Pathog. 2018;14(2):1–23.

97. Ma W, Li S, Ma S, Jia L, Zhang F, Zhang Y, et al. Zika virus causes testis damage and leads to male infertility in mice. Cell. 2016;167(6):1511–24.

98. Clancy CS, Van Wettere AJ, Morrey JD, Julander JG. Zika Virus Associated pathology and Antigen presence in the testicle in the Absence of sexual transmission During subacute to Chronic Infection in a Mouse Model. Sci Rep. 2019;9(1):8325.

99. Demir A, Türker P, Önol FF, Sirvanci S, Findik A, Tarcan T. Effect of experimentally induced Escherichia coli epididymoLorchitis and ciprofloxacin treatment on rat spermatogenesis. Int J Urol. 2007;14(3):268–72.

100. Deng Y-Q, Zhang N-N, Li C-F, Tian M, Hao J-N, Xie X-P, et al. Adenosine analog NITD008 is a potent inhibitor of Zika virus. In: Open forum infectious diseases. Oxford University Press; 2016. p. ofw175.

101. Li C, Deng YQ, Wang S, Ma F, Aliyari R, Huang XY, et al. 25-Hydroxycholesterol Protects Host against Zika Virus Infection and Its Associated Microcephaly in a Mouse Model. Immunity. 2017;46(3):446–56.

102. Grant A, Ponia SS, Tripathi S, Balasubramaniam V, Miorin L, Sourisseau M, et al. Zika virus targets human STAT2 to inhibit type I interferon signaling. Cell Host Microbe. 2016;19(6):882–90.

103. Kumar A, Hou S, Airo AM, Limonta D, Mancinelli V, Branton W, et al. Zika virus inhibits typeLI interferon production and downstream signaling. EMBO Rep. 2016;17(12):1766–75.

104. Wu Y, Liu Q, Zhou J, Xie W, Chen C, Wang Z, et al. Zika virus evades interferon-mediated antiviral response through the co-operation of multiple nonstructural proteins in vitro. Cell Discov. 2017;3:17006.

105. Smith DR, Hollidge B, Daye S, Zeng X, Blancett C, Kuszpit K, et al. Neuropathogenesis of Zika Virus in a Highly Susceptible Immunocompetent Mouse Model after Antibody Blockade of Type I Interferon. PLoS Negl Trop Dis. 2017;11(1):1–22.

106. Xia H, Luo H, Shan C, Muruato AE, Nunes BTD, Medeiros DBA, et al. An evolutionary NS1 mutation enhances Zika virus evasion of host interferon induction. Nat Commun. 2018;9(1).

107. Chen J, Yang YF, Yang Y, Zou P, Chen J, He Y, et al. AXL promotes Zika virus infection in astrocytes by antagonizing type i interferon signalling. Nat Microbiol. 2018;3(3):302–9.

108. McDonald EM, Duggal NK, Delorey MJ, Oksanish J, Ritter JM, Brault AC. Duration of seminal Zika viral RNA shedding in immunocompetent mice inoculated with Asian and African genotype viruses. Virology. 2019;

109. Gorman MJ, Caine EA, Zaitsev K, Begley MC, Weger-Lucarelli J, Uccellini MB, et al. An Immunocompetent Mouse Model of Zika Virus Infection. Cell Host Microbe. 2018;23(5):672-685.e6.

110. Salazar V, Jagger BW, Mongkolsapaya J, Burgomaster KE, Dejnirattisai W, Winkler ES, et al. Dengue and Zika virus cross-reactive human monoclonal antibodies protect against Spondweni virus infection and pathogenesis in mice. Cell Rep [Internet]. 2019;26(6):1585-1597.e4. Available from: https://doi.org/10.1016/j.celrep.2019.01.052

111. Mansuy JM, Dutertre M, Mengelle C, Fourcade C, Marchou B, Delobel P, et al. Zika virus: High infectious viral load in semen, a new sexually transmitted pathogen? Lancet Infect Dis. 2016;16(4):405.

112. D’Ortenzio E, Matheron S, Yazdanpanah Y. Evidence of sexual transmission of Zika Virus. N Engl J Med. 2016;374(22):2195–8.

113. Sheridan MA, Yunusov D, Balaraman V, Alexenko AP, Yabe S, Verjovski-Almeida S, et al. Vulnerability of primitive human placental trophoblast to Zika virus. Proc Natl Acad Sci. 2017;114(9):E1587–96.

114. Wu KY, Zuo GL, Li XF, Ye Q, Deng YQ, Huang XY, et al. Vertical transmission of Zika virus targeting the radial glial cells affects cortex development of offspring mice. Cell Res. 2016;26(6):645–54.

115. Lazear HM, Govero J, Smith AM, Platt DJ, Fernandez E, Miner JJ, et al. A mouse model of Zika virus pathogenesis. Cell Host Microbe. 2016;19(5):720–30.

116. Duggal NK, Ritter JM, Pestorius SE, Zaki SR, Davis BS, Chang G-JJ, et al. Frequent Zika virus sexual transmission and prolonged viral RNA shedding in an immunodeficient mouse model. Cell Rep. 2017;18(7):1751–60.

